# Model-supported patient stratification using multi-objective synergy optimization in combination therapy

**DOI:** 10.64898/2026.05.04.722754

**Authors:** Jana L. Gevertz, Irina Kareva

## Abstract

The challenge of stratifying patients for combination therapy is both technically demanding and clinically crucial. In previous work, we introduced a multi-objective optimization framework for identifying optimally synergistic combination protocols that are robust to competing definitions of additivity. This manuscript extends this methodology to quantify how inter-individual variability in drug sensitivity influences the combination doses that optimally balance the competing objectives of synergy of efficacy and synergy of potency (a proxy measure of toxicity). For this methodology, we introduce a voxel-based stratification approach to characterize individuals (model parameterizations) into subgroups based on sensitivity to each drug as a monotherapy and in combination. As a case study, we apply the method to a preclinical dataset of murine response to the combination of an immune checkpoint inhibitor and an antiangiogenic agent. We demonstrate that the algorithm can quantify how the robustly optimal combination therapies vary across different treatment response subgroups and how the algorithm can identify subpopulations for which no meaningfully efficacious combination exists. As applying the methodology requires knowledge of specific parameter values for which measurable biomarkers may be unavailable, we also propose an initiation protocol that permits identification of the parameters necessary to place an individual in a subgroup. This methodology is a step in the direction of determining the right combination therapy for a subgroup and finding the right subgroup for an existing therapy.

## 1. INTRODUCTION

Clinical trials are designed to determine the efficacy and safety of new drugs for the average patient. However, it is now well-established that treatment benefits and risks are not uniform across the patient population (Kent et al. 2020), and that conclusions about the average patient do not necessarily apply to an individual (Luo et al. 2022). In trials where a drug shows only modest benefit in the full population, meaningful effects may still exist within a patient subgroup (Tanniou et al. 2016). The stratification of patients into subgroups, typically done using a predictive biomarker, can thus effectively “rescue” a compound that might otherwise appear ineffective or intolerable.

There are many recent examples demonstrating how the use of predictive biomarkers can help ensure that therapies reach the patients that are most likely to respond. For instance, the epidermal growth factor receptor (EGFR) tyrosine-kinase inhibitor gefitinib (brand name Iressa) initially gained accelerated approval from the FDA in 2003 for advanced non-small cell lung cancer (NSCLC). However, a crucial Phase III study comparing gefitinib to a placebo in the Iressa Survival Evaluation in Lung Cancer (ISEL) trial failed to demonstrate a significant improvement in overall survival, leading to restricted use of gefitinib in 2005 (Karachaliou et al. 2019). It was only after retrospective biomarker analyses revealed that responses were limited to tumors harboring activating EGFR mutations (exon 19 deletions or L858R) that the benefit of treatment became evident. Subsequent trials in EGFR-mutant cohorts then demonstrated significantly improved response rates (Costa et al. 2007). This led to gefitinib receiving renewed approval from the FDA in 2015 as a first-line treatment for metastatic NSCLC patients whose tumors harbor these specific EGFR mutations (Kazandjian et al. 2016).

There are many other examples within oncology. Initial trials of trastuzumab in unselected breast cancer patients showed negligible benefit until human epidermal growth factor receptor 2 (HER2) overexpression or amplification was incorporated into the inclusion criteria (Xia et al. 2023). Similarly, early inhibitors of the poly (ADP-ribose) polymerase (PARP) enzyme in unselected ovarian and breast cancer showed minimal activity. However, when restricted to germline BRCA1/2 mutation carriers, monotherapy with the PARP inhibitor olaparib was approved for BRCA-mutant ovarian cancer after multiple prior lines of therapy (Domchek et al. 2016). Another example is that of the BRAF inhibitor vemurafenib, which failed to show benefit in unselected melanoma. However, in the BRIM-3 trial selecting only BRAF V600E-mutant tumors, it improved progression-free and overall survival (Chapman et al. 2017). For this genetically defined subgroup, combination with cobimetinib (a MEK inhibitor) further extended progression free survival and overall survival.

These examples illustrate that, without prospective biomarker stratification, real signals of efficacy can become diluted in the general population. As such, treatment protocols must be coupled with the right patient population. Mathematical modeling approaches have been increasingly used to facilitate this process (Scibilia et al. 2025). An example is the use of a mathematical model of adaptive therapy for abiraterone in metastatic castrate-resistant prostate cancer, where the treatment was stopped when prostate specific antigen (PSA) fell to below 50% of baseline and restarted when it returned to baseline. The authors predicted longest disease control for those individuals whose PSA rebounded slowly, while fast-rebound patients progressed earlier (Brady-Nicholls et al. 2021). In another example, Avanzini et al. (Avanzini et al. 2020) analyzed whether negative circulating tumor DNA truly implies minimal disease. The authors predicted that high-shedding patients were reliably “seen” early (actionable negatives), whereas low-shedding/small tumors risked false negatives and needed imaging-heavy follow-up. In (Gevertz & Kareva 2025), we used a soluble biomarker, B-cell maturation antigen and a virtual population analysis to propose an approach for identifying subgroups of multiple myeloma patients that may benefit from dose adjustments for bispecific T cell engager teclistamab.

Crucially, part of what determines success of therapy is not just efficacy but toxicity. Chemotherapy-induced toxicities can lead many patients to receive lower doses than is indicated (Nielson et al. 2021). Treatment discontinuation due to adverse events has also been reported for colon cancer (Boakye et al. 2021), osteosarcoma (Spreafico et al. 2024), as well as in immunotherapy (Shin et al. 2024), often compromising efficacy. A successful therapeutic approach must balance both efficacy and toxicity, something that may be particularly challenging to execute for a combination therapy, where multiple toxicity profiles need to be monitored in addition to efficacy.

One approach to address this issue is by using the notion of Pareto optimality for balancing the competing objectives of efficacy and toxicity in combination therapy. A Pareto optimal dose is one where one objective cannot be improved without compromising the other. In this work, we apply this framework to an updated mathematical model of combination therapy with the checkpoint inhibitor pembrolizumab and the anti-angiogenic agent bevacizumab. We introduce treatment response heterogeneity into the model and showcase the differences in Pareto optimal dose selection for individuals with different sensitivities to the drugs. We also use this framework to identify the subgroups for which a pre-determined combination protocol optimally balances synergy of efficacy and potency. We conclude with a proposed method to identify individual parameters that cannot be measured directly. These parameters can then be thought of as a mathematical biomarker that permits patient stratification.

## 2. METHODS

Here, we introduce an expansion of a previously developed framework for balancing synergy of efficacy and potency (a proxy measurement of toxicity) of combination therapies called MOOCS-DS (Multi-Objective Optimization of Combination Synergy – Dose Selection) (Gevertz & Kareva 2023). As a case study for illustrating the applicability of the method, we consider a model of the combination of the PD-1 checkpoint inhibitor pembrolizumab, and the anti-angiogenic agent bevacizumab calibrated to preclinical data. We use the model to demonstrate how the framework for balancing combination efficacy and toxicity can be applied to study the heterogenous response to combination therapy. All simulations are conducted in MATLAB and the code is available at https://github.com/jgevertz/MOOCS-DS/tree/main/Variability.

### 2.1 Pareto Optimization of Synergy via an Expanded MOOCS-DS Algorithm

The biomedical community has long sought to identify drug combinations for which the combined effect is synergistic, defined as being greater than additive. However, lack of consensus on the definition of additivity has complicated this goal (Gevertz & Kareva 2023)(Vlot et al. 2019). It is common for the same combination to be classified as synergistic by one additivity definition and antagonistic (when the outcome is less than expected in the additive case) by another, making these classifications difficult to interpret. The definitions of additivity, and thus synergy, generally fall into one of two categories: those focused on the output of the treatment (synergy of efficacy, SoE) and those focused on the input of the treatment (synergy of potency, SoP).

In previous work (Gevertz & Kareva 2023), we developed the MOOCS-DS algorithm to identify synergistically optimal combination therapies that do not sensitively depend on how additivity has been defined (see Supplementary Methods for details). In particular, rather than focusing on just one form of synergy (efficacy or potency), we choose to simultaneously optimize both types of synergy. Further, rather relying on a single definition of synergy of efficacy and potency, we consider a set of definitions for each. For synergy of efficacy we consider additivity as defined using both Bliss and Highest Single Agent (HSA). For synergy of potency, we consider additivity as defined using both Loewe and Lowest Single Dose (LSD). Details of these definitions are provided in the Supplementary Methods Section.

The MOOCS-DS algorithm performs Pareto optimization in *N*_*spaces*_ = 4 SoP-SoE criterion spaces: Loewe-Bliss, Loewe-HSA, LSD-Bliss, LSD-HSA. Pareto optimal doses are identified in each of the four spaces and all doses identified as Pareto optimal are assigned a score from 0-4. A score of 0 indicates that the dose is never identified as Pareto optimal, whereas a score of 4 indicates that the dose is Pareto optimal in all *N*_*spaces*_ SoP-SoE criterion spaces. Herein, we expand the original MOOCS-DS algorithm in two key ways.

In the original implementation, doses that would never be used in practice can be identified as Pareto optimal. This can occur because Pareto optimality only looks to balance synergy of potency and efficacy, but it does not impose any minimal expectation on efficacy (or any maximally acceptable toxicity). Given that Pareto optimization results in a set of solutions (the Pareto front) and not a single solution, one option is to simply ignore any Pareto optimal solutions that do not meet minimum standards of efficacy (or potency).

However, since Pareto optimality is assessed relative to existing solutions, a more mathematically sound approach is to first impose any constraints on efficacy (and/or toxicity) and then perform a constrained Pareto optimization. Therefore, the updated version of MOOCS-DS permits constraints on efficacy to be imposed prior to performing dose optimization. Specifically, one can set a minimally sufficient efficacy threshold for the combination, and any doses that do not reach that threshold are excluded from analysis. Setting the threshold to zero would recover the original implementation of MOOCS-DS. Unless otherwise specified, all results shown in this paper use an efficacy threshold of 0.6. This means that if a combination therapy does not achieve a 60% reduction in tumor size relative to control, the combination dose is removed from consideration prior to performing the Pareto optimization. Notably, in this implementation of the algorithm no explicit constraints on toxicity were added. Instead, we assume that a toxicity constraint is indirectly implemented by imposing a maximum allowable dose for each drug.

Additionally, the expanded MOOCS-DS algorithm can now identify solutions we call “near Pareto optimal”, rather than only Pareto optimal solutions. As illustrated in Supplementary Figure S1, there can be points in the criterion space that have objective function values that are very similar to the values on the Pareto front, yet these points are classified as non-optimal because another dose balances the two criteria slightly better. Given that for practical purposes there is no difference between a dose that falls precisely on the Pareto front, and a dose that falls very close to the front, the MOOCS-DS algorithm now offers the option to identify both Pareto optimal doses and near Pareto optimal doses that fall within some fixed (small) distance from the Pareto front.

To identify near Pareto points, we must first define a distance metric from the Pareto front. This metric must account for the fact that area of the criterion space that the points are mapped into varies depending on the additivity metrics used. To normalize across different criterion spaces, we identify the point in criterion space closest to the origin, and the point further from the origin. We then compute the Euclidean distance, *L*^2^, between these points (*pdist2* in MATLAB). This distance is then used as a single metric that defines the “extent” of the region. The Pareto front is next represented mathematically using finely resolved piecewise linear interpolation (computed using *interp1* in MATLAB). The Euclidean distance *d*_*i*_ of each non-Pareto optimal point to the Pareto front can then be calculated. A point is defined as near Pareto optimal provided *d*_*i*_ ≤ ∈*L*^2^, where 0 ≤ ∈ ≤ 1 controls how strictly we define a near Pareto optimal point. Setting ∈ to zero results in the algorithm only identifying Pareto optimal solutions as in the original implementation, whereas setting ∈ to one results in every point in the space being classified as near Pareto optimal. In practice, ∈ should be quite small. We use ∈ = 0.02.

MOOCS-DS was designed to be robust to uncertainties in how additivity (and thus synergy) is defined. In this work, we use the expanded version of the algorithm to also test the robustness of optimally synergistic combinations to population-level heterogeneity. In practice, this means we will explore Pareto optimal solutions across parameter space, as detailed in the Parameter Variability Subsection.

### 2.2 Model description

In (Gevertz & Kareva 2023), we developed a model to capture the impact on tumor burden of a combination of two existing drugs, a checkpoint inhibitor pembrolizumab and an anti-angiogenic agent bevacizumab, administered in two mouse models of non-small cell lung cancer (NSCLC); here we focus on the results from the more sensitive p53-deficient H1299 cell line. The animals received either: 1) 10 mg/kg of pembrolizumab starting on day 17, then given every 3 days (Q3D) until five doses have been administered, 2) 1 mg/kg of bevacizumab starting on day 14, then given every 3 days until six doses have been administered, or 3) a combination of these two drugs using the same protocol.

A standard pharmacokinetic (PK)-tumor growth inhibition (TGI) model was used for both drugs given as monotherapy or in combination as follows:

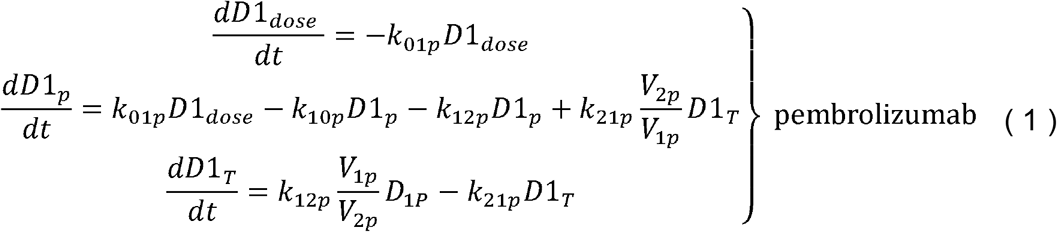

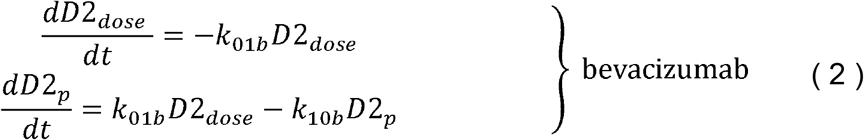

Drug PK for bevacizumab were best described by a standard one-compartment model, parameterized using data from Lin et al. (Lin et al. 1999). Drug PK for pembrolizumab was best described by a standard two-compartment model parametrized with data from Lindauer et al. (Lindauer et al. 2017). For both models, it is assumed that the drug is administered intravenously into the central (plasma) compartment at a rate constant *k*_01_ and that the drug can be cleared from the plasma compartment at a rate constant *k*_10_. Pembrolizumab can also distribute into the peripheral compartment at a rate constant *k*_12*p*_ and return into the central compartment at a rate constant *k*_21p_. Finally, it is assumed that the tumor grows logistically and can be killed as a function of concentration of either of the drugs.

Here, we have updated the tumor growth model, changing the terms describing drug sensitivity to each drug from the original mass action terms (Gevertz & Kareva 2023) to Michaelis-Menten saturation terms. We have also removed the compartments that represent the transition of cancer cells through multiple stages before eventually dying off, as these transitional compartments proved unnecessary to adequately fit the data. The resulting equation that describes tumor (*x*) dynamics is as follows:

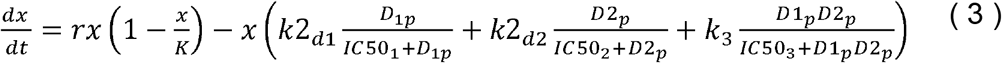

The results are shown in Figure 1; the updated parameter values and nonzero initial conditions are summarized in Table ***1***.

**Figure 1.**
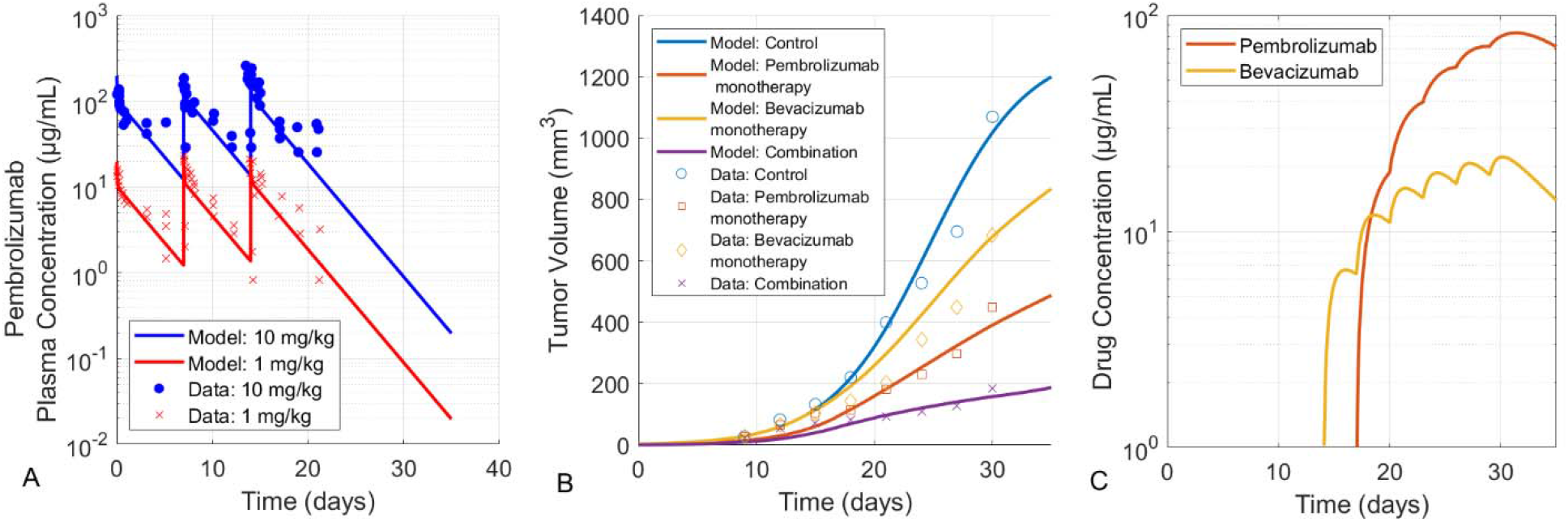
(A) Pembrolizumab PK, model vs data; data were digitized from (Lindauer et al. 2017). PK parameters for bevacizumab remain unchanged compared to (Gevertz & Kareva 2023) and therefore no updated PK plots are shown. (B) Predicted tumor growth inhibition curves for the protocols, described in (Qiao et al. 2023). (C) Drug concentration profiles (on a log scale) as simulated using the proposed PK models (Table 1. Baseline parameters used in System (1)-(3). The drug profiles are simulated for the same combination protocol used by (Qiao et al. 2023).

**Table 1.** Baseline parameters used in System (1)-(3).

| Parameter | Description | Value | Units | Ref. |
| --- | --- | --- | --- | --- |
| **Pembrolizumab** |  |  |  |  |
| $V_{1p}$ | Volume of distribution, central compartment | 59.6 | ml/kg | Calibrated to data in (Lindauer et al. 2017) |
| $V_{2p}$ | Volume of distribution, peripheral compartment | 44.45 | ml/kg | |
| $Cl_{1p}$ | Clearance, central compartment | 29.28 | ml/kg/day | |
| $Cl_{2p}$ | Clearance, peripheral compartment | 450 | ml/kg/day | |
| $k_{10p}$ | Rate of drug clearance from the central compartment for drug | $Cl_{1p}/V_{1p}$ | 1/day | |
| $k_{12p}$ | Rate constant for drug distribution from central to peripheral compartment | $Cl_{2p}/V_{1p}$ | 1/day | |
| $k_{21p}$ | Rate constant for drug distribution from peripheral to central compartment | $Cl_{2p}/V_{2p}$ | 1/day | |
| $k_{01p}$ | Drug absorption rate for subcutaneous (sc) administration | 0.11 | 1/day | |
| **Bevacizumab** |  |  |  |  |
| $V_{1b}$ | Volume of distribution, central compartment | 119 | ml/kg | Calibrated to data in (Lin et al. |
| $V_{2b}$ | Volume of distribution, peripheral compartment | 13.6 | ml/kg/day | |
| $k_{10b}$ | Rate of drug clearance from the central compartment for drug | $Cl_{1b}/V_{2b}$ | 1/day | 1999) |
| $k_{01b}$ | Drug absorption rate for subcutaneous (sc) administration | 1.3 | 1/day | |
| Tumor (H1299) |  |  |  |  |
| K | Tumor carrying capacity | 1300 | $\text{mm}^3$ | Calibrated to data reported in (Qiao et al. 2023) |
| r | Intrinsic rate of tumor growth | 0.24 | 1/day |  |
| $k_{2d1}$ | Rate of tumor elimination by pembrolizumab as monotherapy | 0.15 | 1/day | |
| $k_{2d2}$ | Rate of tumor elimination by bevacizumab as monotherapy | 0.09 | 1/day | |
| $k_3$ | Rate of tumor kill by combination of bevacizumab and pembrolizumab | 0.002 | 1/day | |
| $IC50_1$ | Half-maximal concentration of pembrolizumab as monotherapy | 25 | nM | |
| $IC50_2$ | Half-maximal concentration of bevacizumab as monotherapy | 10 | nM | |
| $IC50_3$ | Half-maximal concentration of pembrolizumab combined with bevacizumab | 15 | nM | |
| $x(0)$ | Initial tumor volume | **Control:** 3.5 on day 0; 133 on day 15<br>**Drug 1 monotherapy** : 1.75 on day 0, 105 on day 15<br>**Drug 2 monotherapy** : 3.5 on day 0, 105 on day 15<br>**Combination** : 1.167 on day 0, 71 on day 15 | $\text{mm}^3$ | |

### 2.3 Parameter Variability

To assess the impact of inter-individual heterogeneity in response to combination therapy, we introduce variability into the drug response parameters. We focus on the tumor kill rates associated with each monotherapy (*k*2_*d*1_, *k*2_*d*2_) and the kill rate associated with the combination (*k*_3_). All PK parameters, along with the intrinsic tumor growth rate and the IC50 values, were fixed at their nominal value (Table ***1***). The approach can, however, be expanded to include additional parameters as needed.

As no inter-individual variability data were available in (Qiao et al. 2023), we chose to define the upper bound on each parameter as the value that results in the final tumor volume being 50% smaller than the volume at the nominal parameter value, and the lower bound as the value that results in the predicted final tumor volume being 50% larger than predicted by the nominal parametrization (Figure 2A-C). When this criterion cannot be satisfied, the lower bound on the parameter is set to be three orders of magnitude smaller than the nominal value. (The criterion was always able to be satisfied for the upper bound on the parameter.) Parameter bounds determined in this way create a parameter cuboid (Figure 2D).

**Figure 2.**
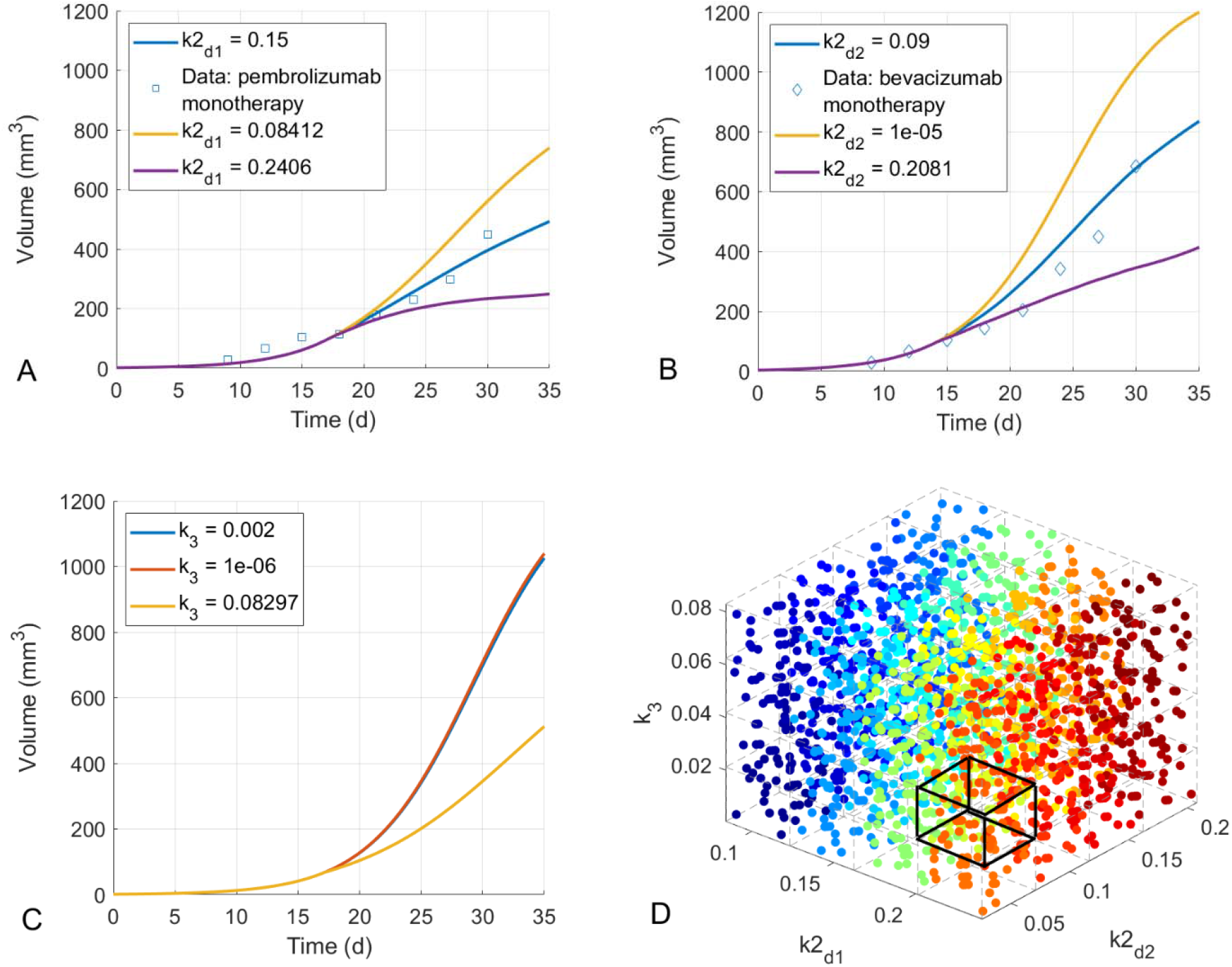
Parameter values associated with 50% change in end tumor volume for (A) pembrolizumab monotherapy, (B) bevacizumab monotherapy, (C) the effect of only the combination kill term when administering the combination therapy. We note that it was not possible to achieve a 50% increase final tumor volume in the case of bevacizumab monotherapy (varying *k*2_*d*2_) and in the case of the combination therapy (setting *k*2_*d*1_ = *k*2_*d*2_ = 0 and varying *k*_3_), as a 50% increase results in a tumor larger than the control volume. In those cases, a lower bound on the parameter was set to be three orders of magnitude smaller than the nominal value. (D) The full parameter cuboid, decomposed into 64 voxels. Precise boundaries for each voxel are described in the Online Supplementary Material (Supplementary File 1). Black outline highlights a representative voxel. Each point represents a sampled parameter set within the cuboid, with points in the same voxel shown using the same color.

Next, we subdivide each dimension of the cuboid (*k*2_*d*1_, *k*2_*d*2_, *k*_3_) into four equal-sized subregions. This can be thought of as classifying treatment response into one of four categories: minimal response, mild response, moderate response, and strong response. We note that these labels represent relative treatment response along a single axis in parameter space, but not an absolute response to the combination protocol in the full three-dimensional parameter space (see Figure 2D). While a different number of subregions could be chosen, four allows for sufficient variability in response to monotherapy and combination therapy while limiting computational complexity. This creates 64 possible response categories (four for each of pembrolizumab monotherapy, bevacizumab monotherapy, and the combination), which we will call a parameter voxel. The parameter cuboid with a representative voxel is shown in Figure 2D.

Our goal is to understand the response of “individuals” that fall within each parameter voxel. In this context, an individual is defined as a parameterization of the mathematical model. To this end, we randomly sample, using Sobol sequences (MATLAB command *sobolset*), *N*_*samples*_ = 30 parameterizations from each voxel. MOOCS-DS is then executed for each of voxel’s 30 parameterizations, resulting in 30×64 = 1920 runs of MOOCS-DS. We then compute the voxel-specific statistics regarding (near) Pareto optimal combinations. This approach ensures that the individuals within each voxel are similar enough, so that a recommendation can be made with confidence that a combination is optimally synergistic for this subgroup. As an example, a voxel could represent minimal responders to pembrolizumab monotherapy, moderate responders to bevacizumab monotherapy, and mild responders to combination effects.

The main metric we will use to evaluate the robustness of a Pareto optimal prediction across definitions of synergy and intra-voxel individual variability will be what we call the *Pareto Optimality Score, 𝒫* :

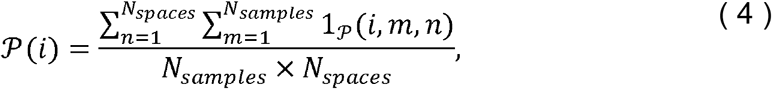

where *N*_*samples*_ is the number of parameterizations sampled per voxel (*N*_*samples*_ = 30 here), *N*_*spaces*_ is the number of criterion spaces in which Pareto optimization was performed (*N*_*spaces*_ = 4 here) and 1_𝒫_ (*m*,*n* is the indicator function defined as:

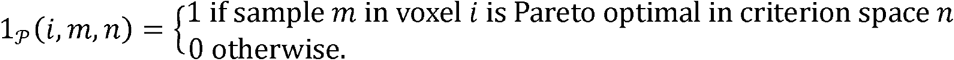

The voxel-specific Pareto Optimality Scores can then be used to answer two types of questions: 1) If we know to which voxel an individual belongs, what would be a robust Pareto optimal combination dose for this individual? 2) Alternatively, if a combination dose is fixed (i.e., a protocol has been established and cannot be easily modified), for individuals belonging to which voxel would this dose be robustly Pareto optimal? And for both cases, we also ask: if we do not have a measurable biomarker, how can individuals be stratified to determine which voxel they belong to?

## 3. RESULTS

First, we highlight the importance of incorporating inter-individual variability on the predicted Pareto optimal combination dose. Recall that in the previous work (Gevertz & Kareva 2023), we analyzed a homogeneous population, where the sensitivity to both monotherapies (*k*2_*d*1_and *k*2_*d*2_) and to combination therapy (*k*_3_) were fixed. However, as shown in Figure 3, Pareto optimal combination regimens can be very different depending on individual sensitivities to the monotherapies. In what follows, for brevity we will adopt the following notation: a dose of, for instance, 10 mg/kg of pembrolizumab with 1 mg/kg of bevacizumab achieving a hypothetical TGI of 60% will be denoted as P10/B1/TGI60.

**Figure 3.**
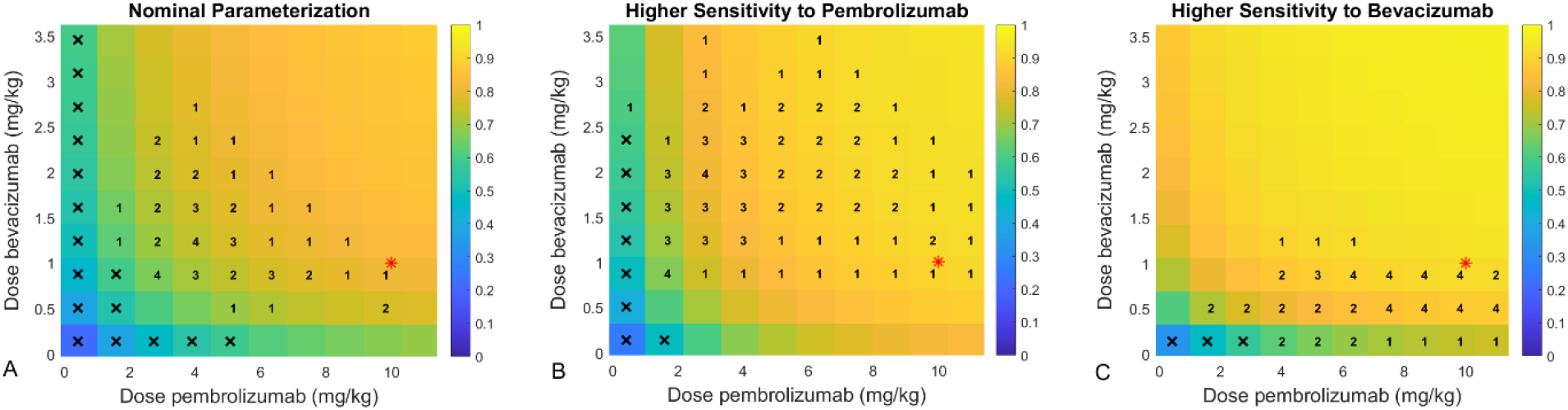
Representative Pareto optimal doses for different individuals. The color in the heatmap indicates the predicted tumor growth inhibition (TGI) relative to the control for the combination dose, with 1 corresponding to 100% TGI. The number indicates the number of criterion spaces (of the four considered) for which a dose is Pareto optimal. The red star shows the dose used in experiments. (A) Nominal parameters: *k*2_*d*1_ = 0.15, *k*2_*d*2_ = 0.09, *k*_3_ =0.002. (B) Parametrization for an individual with a higher sensitivity to pembrolizumab with *k*2_*d*1_ =0.22, *k*2_*d*2_ =0.09, *k*_3_ =0.002. (C) Parametrization for an individual with a higher sensitivity to bevacizumab with *k*2_*d*1_= 0.15, *k*2_*d*2_ = 0.18,*k*_3_ =0.002. An ‘x’ indicates that the combination dose did not meet the efficacy threshold (TGI ≥ 0.6) to be included in the set of doses over which the Pareto optimization is performed.

For instance, in Figure 3A, we show the MOOCS-DS output for the nominal parametrization (reported in Table ***1***) and compare that to the originally tested experimental dose of 10 mg/kg of pembrolizumab and 1 mg/kg of bevacizumab (Qiao et al. 2023), or in our notation P10/B1, denoted by a red star. The combination tested in the dose sweep closest to the experimental protocol is identified as Pareto optimal in only a single criterion space (denoted by a ‘1’ in Figure 3A), indicating a high sensitivity to how synergy of efficacy and potency are defined. The Pareto optimal combinations that were robust to the additivity definitions (denoted by a ‘4’ in Figure 3A) use a much lower dose of pembrolizumab than the experiment protocol (P2.7/B0.9, or P3.9/B1.2).

If we consider an individual with a higher sensitivity to pembrolizumab (Figure 3B), an interesting phenomenon occurs. While the most robust Pareto optimal combinations are neighboring doses for the nominal parameterization (that is, they are nearest neighbors in the dosing grid), the same is not true when the sensitivity to pembrolizumab is increased. The most robust Pareto optimal doses are P1.6/B0.9/TGI67 or P2.7/B2.0/TGI0.81. The protocols that are “in between” these are mostly classified with a ‘3’ in Figure 3B, which indicates strong (though not unanimous) support across multi-objective synergy spaces. This can be interpreted as having a larger dose range for which this individual has a Pareto optimal response. Though, each dose does correspond to a different point on the Pareto front, meaning they represent distinct ways to balance synergy of efficacy and potency.

As another illustrative example, in Figure 3C we show the Pareto optimal doses for an individual with a higher sensitivity to bevacizumab. In this instance, a set of seven combination doses are identified as robustly Pareto optimal. These have pembrolizumab administered in the range of 6.2 – 9.8 mg/kg and bevacizumab 0.5 – 0.9 mg/kg. While the exact experimental dose was not tested due to how dosing space gets discretized when performing the protocol sweep, it is interesting to note that the experimental dose is very close to these robustly Pareto optimal doses. This is in stark comparison with the nominal parameterization and individuals with a higher sensitivity to pembrolizumab.

Taken together, these results highlight that the Pareto optimal combination therapy is sensitive to the treatment-related parameters. While such heterogeneity is to be expected, it generally gets averaged out when recommending a population-level protocol. Such a population-level approach is viable if the range of possible parameter values is sufficiently narrow, or if treatment response is insensitive to parameter variability. In these cases, deviations from the mean do not dramatically change the predicted optimal combination regimen for the population. However, substantial inter-individual variability, such as differences in drug sensitivity that are common in cancer treatment, limits the usefulness of an average dose prediction.

### 3.1 Finding a Pareto optimal combination dose for a known voxel

We now turn to the following question: if we know an individual’s sensitivity to both monotherapies and the combination therapy (that is, we know which voxel of the parameter cuboid they fall in), can we use this to determine a Pareto optimal combination therapy? To address this question, we ran MOOCS-DS for each individual (randomly sampled parameterization) within each voxel, recording the likelihood of each dose being Pareto optimal (i.e., balancing out toxicity and efficacy) across four multi-objective criterion spaces. A representative example from voxel 49, where individuals have a strong response to pembrolizumab monotherapy, a minimal response to bevacizumab monotherapy, and a minimal response to the combination therapy, is shown in Figure 4.

**Figure 4.**
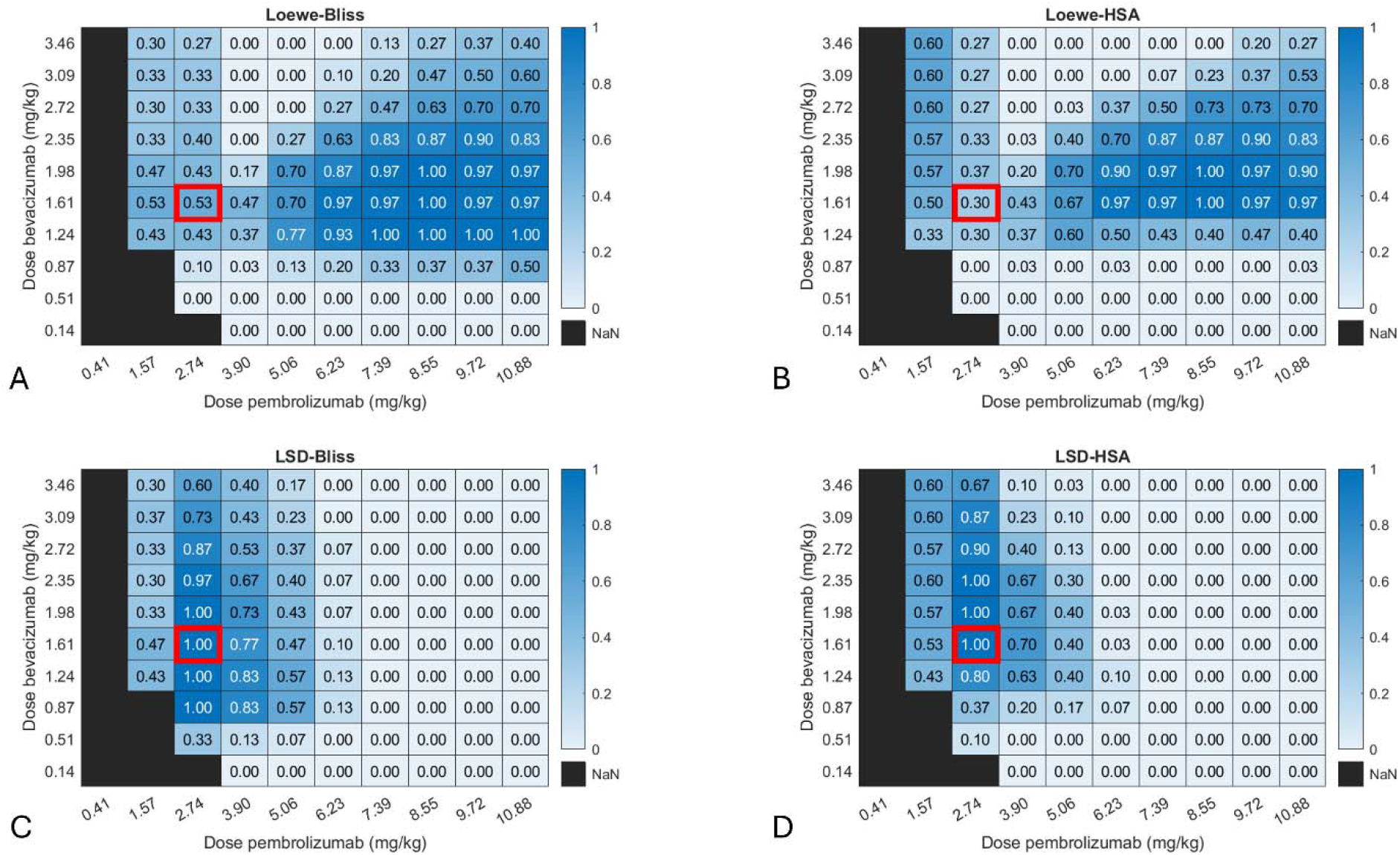
The four multi-objective criterion spaces and the predicted probability of a dose being Pareto optimal for parameters in voxel 49 (, ,). The four multi-objective criterion spaces are: (A) Loewe-Bliss, (B) Loewe-HSA; (C) LSD-Bliss, and (D) LSD-HSA. The red rectangles indicate the combination dose that was identified as having the highest Pareto Optimality Score across all criterion spaces, , for this voxel. A blacked-out cell indicates that the combination dose did not meet the efficacy threshold (TGI) to be included in the set of doses over which the Pareto optimization is performed.

The results in Voxel 49 demonstrate how the Pareto optimal doses vary depending on the metric used to define synergy of efficacy and synergy of potency, consistent with our previous analysis (Gevertz & Kareva 2023). To measure the robustness of a Pareto optimal prediction given this variability, we calculate the *Pareto Optimality Score* defined in Equation (4). The Pareto Optimality Score for voxel 49, , is visualized as a heatmap across dosing space in Figure 5D. In these heatmaps, a value of 1 must mean that every sampled parameterization within the voxel has the dose under consideration as (near) Pareto optimal for all four multi-objective criterion spaces. A value of 0 must mean the dose is never identified as Pareto optimal. Intermediate values are harder to interpret. For instance, a could mean that the dose is Pareto optimal for all sampled parameters, but only in half the multi-objective criterion spaces. Another possible interpretation of is that the dose is Pareto optimal for half of the sampled parameterizations but in all the multi-objective criterion spaces.

**Figure 5.**
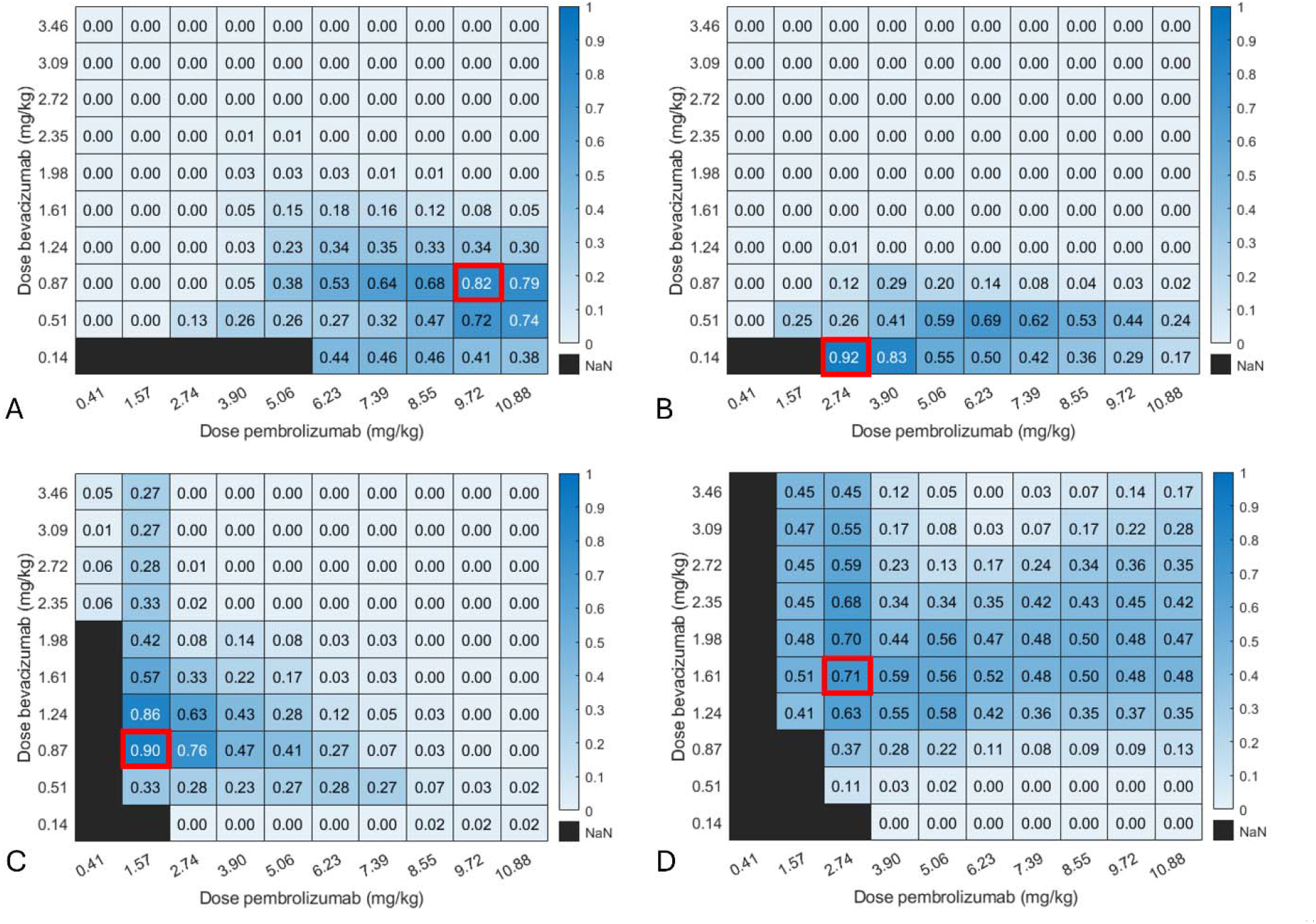
Pareto Optimality Score for four voxels. (A) Output for voxel 13 where individuals have a minimal response to pembrolizumab monotherapy, a strong response to bevacizumab monotherapy, and a minimal response to the combination therapy. (B) Output for voxel 30 where individuals have a mild response to pembrolizumab monotherapy, a strong response to bevacizumab monotherapy, and a mild response to the combination therapy. (C) Output for voxel 35 where individuals have a moderate response to pembrolizumab monotherapy, a minimal response to bevacizumab monotherapy, and a moderate response to the combination therapy. (D) Output for voxel 49 where individuals have a strong response to pembrolizumab monotherapy, a minimal response to bevacizumab monotherapy, and a minimal response to the combination therapy. Doses with highest Pareto optimality scores for each voxel are highlighted. A blacked-out cell indicates that the combination dose did not meet the efficacy threshold (TGI ≥ 0.6) to be included in the set of doses over which the Pareto optimization is performed.

**Figure 6.**
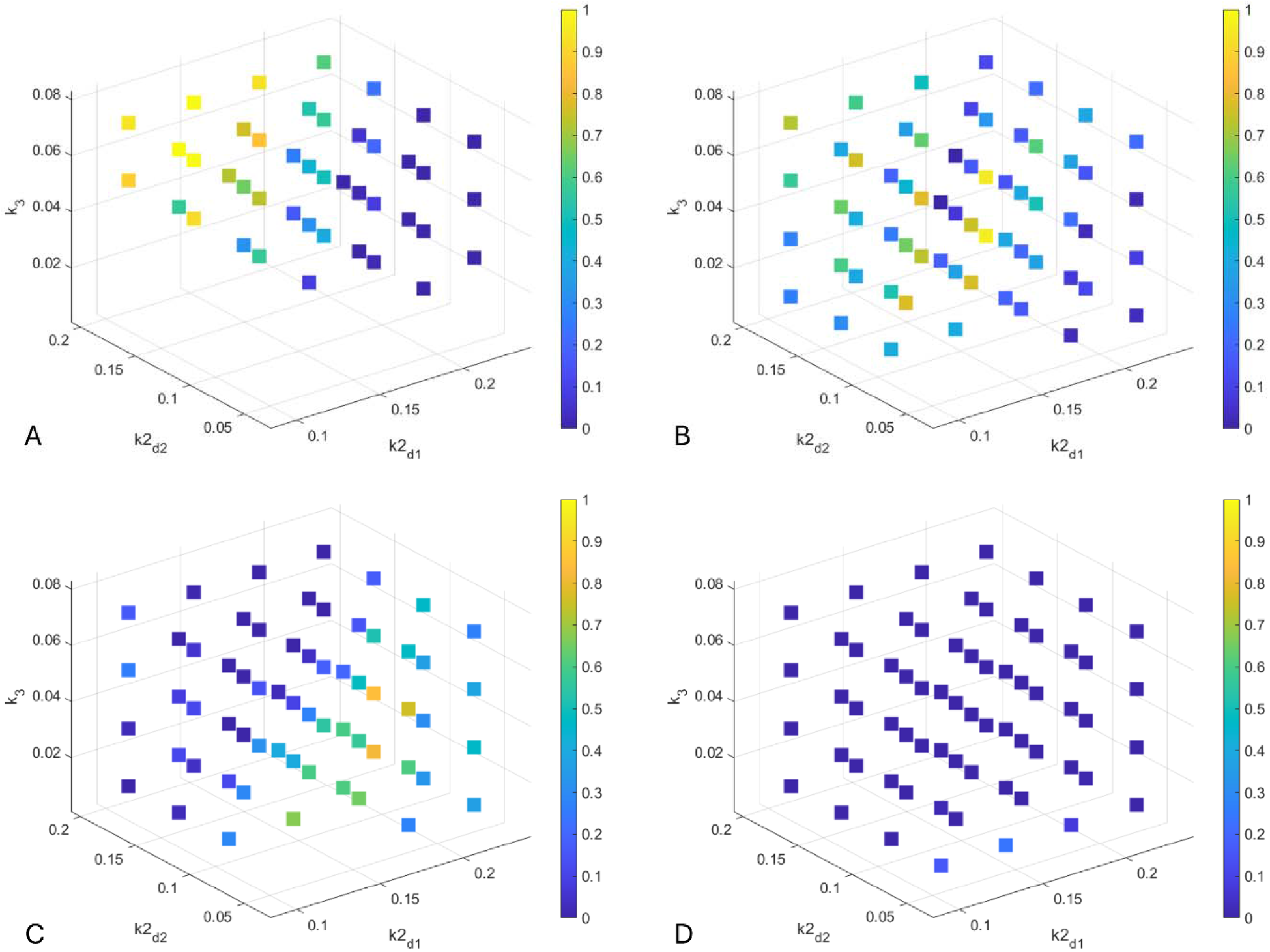
Pareto Optimality Score for all parameter voxels at four representative combination doses. (A) Pembrolizumab dose of 2.7 mg/kg, bevacizumab dose of 0.1 mg/kg; (B) pembrolizumab dose of 3.9 mg/kg, bevacizumab dose of 0.5 mg/kg; (C) pembrolizumab dose of 2.7 mg/kg, bevacizumab dose of 0.9 mg/kg; (D) pembrolizumab dose of 6.2 mg/kg, bevacizumab dose of 3.5 mg/kg. Parameterizations that did not meet the efficacy threshold (TGI ≥0.6) have no value shown.

**Figure 7.**
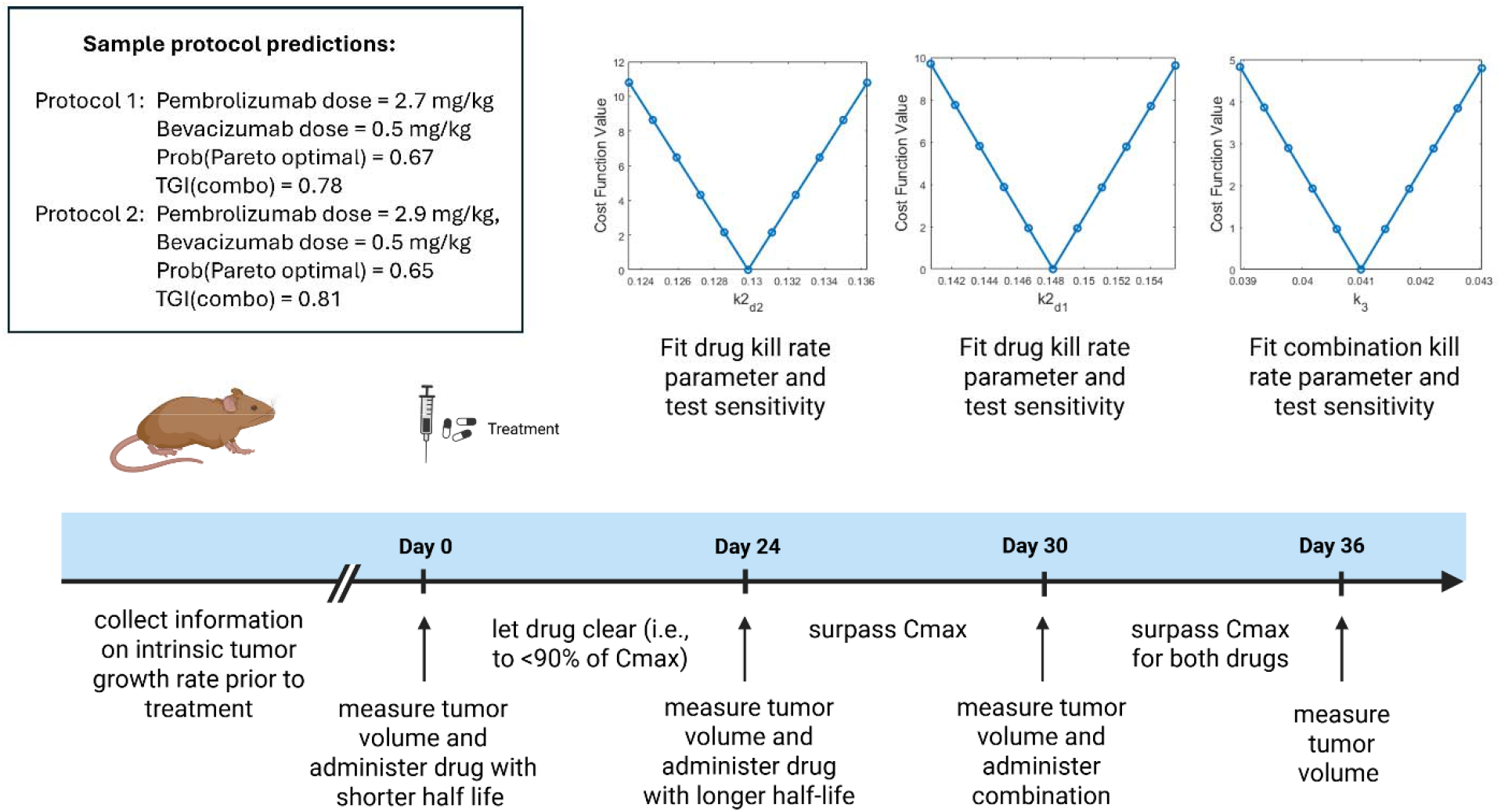
Schematic diagram of the proposed initiation protocol, with sample outputs of how parameter estimates were obtained, and top combination doses for a sample individual with parameters in the identified range.

For individuals in voxel 49, the highest achieved is 0.71, indicating some lack of robustness either within a voxel or across multi-objective criterion spaces. The contribution of each of these factors can be deciphered by looking at the criterion-space specific probabilities (Figure 4). If we focus on the combination with the highest (P2.7/B1.6) and look at each individual criterion space, we see that this dose is Pareto optimal for all sampled parameterizations when LSD is the measure of synergy of potency (Figure 4C,D). However, if Loewe is used instead (Figure 4A,B) only 30% (when paired with HSA) or 53% (when paired with Bliss) of parameterizations within the voxel have this dose as Pareto optimal. Thus, the lack of robustness (distance of 𝒫 from 1 at the dose that maximizes 𝒫) is largely attributable to how synergy of potency is defined. Though, within the criterion spaces that use Loewe additivity, there is also inter-individual variability in whether this dose is (near) Pareto optimal.

Results of this analysis for other representative voxels are shown in Figure 5A, B and C; the relative placement of these voxels within the parameter space is shown in Supplementary Figure S2. Voxel 13 (Figure 5A) represents individuals who have a minimal response to pembrolizumab monotherapy, a strong response to bevacizumab monotherapy, and a minimal response to the combination therapy (minimal/strong/minimal). The dose most likely to be identified as Pareto optimal administers P9.7/B0.9. The Pareto optimality score of 𝒫 = 0.82 indicates more robustness than observed for individuals at the most robust dose in Voxel 49. Notably, for these individuals, the identified protocol is very close to the doses of P10.0/B1.0 that were selected experimentally in (Qiao et al. 2023). Thus, individuals who fall in Voxel 13 are precisely those who we predict will most benefit, in terms of balancing efficacy and potency, from the experimental protocol.

Compare individuals who fall in Voxel 13 to those who fall in Voxel 30 (Figure 5B), which represents those who have a mild response to pembrolizumab, a strong sensitivity to bevacizumab and a mild response to the combination (mild/strong/mild). For these individuals, the highest score achieved is 𝒫 = 0.92 , which is the most robust response we see across the voxels in Figure 5. The dose that was identified as being robustly Pareto optimal for individuals that fall within this voxel gives P2.7/B0.1 per dose. Notably, these doses are much lower than the doses suggested in the original experiment; for the original doses of P10.0/B1.0, the Pareto optimality score for this voxel is near zero. Thus, individuals who fall into this voxel are predicted to be overtreated using the experimental protocol.

A comparison of mild/ strong/ mild individuals who fall in Voxel 30 (Figure 5B) versus the moderate/minimal/moderate individuals who fall in Voxel 35 (Figure 5C) is also quite interesting. The most robust Pareto optimal combination for the Voxel 30 subgroup administers P2.7/B0.1. This same dose for individuals in Voxel 35 is classified as non-optimal in every multi-objective criterion space (𝒫 = 0). Similarly, the most robust Pareto optimal combination for the Voxel 35 subgroup (P1.6/B0.9) has a Pareto optimality score of zero for the Voxel 30 subgroup. This highlights how a combination dose that is Pareto optimal for one subgroup can perform very poorly in a different subgroup.

We note that while the algorithm is initialized using 100 possible combination doses, the number that satisfy the TGI constraint imposed prior to implementing the multi-objective optimization routine varies across parameter space. The weakest responders fall in Voxel 1 – their parameters are such that response to each monotherapy and the combination therapy is minimal. For these individuals, the larger the TGI threshold set, the smaller the set of feasible doses becomes (Supplementary Table S1). Focusing on this parameter voxel, the TGI threshold utilized herein (0.6 tumor growth inhibition relative to control) was chosen because it permits half the doses to be included in the feasible set. Using a TGI threshold much beyond this quickly decreases the number of feasible doses. In fact, a TGI threshold of 0.75 or higher results in an empty feasible set (Supplementary Table S1, Supplementary Figure S3). Thus, it is important to be aware that even the dose(s) identified as Pareto optimal (in terms of balancing efficacy and potency) may not achieve a sufficiently high efficacy for clinical use.

A video showing the results for all 64 voxels can be found in the Online Supplementary Materials.

### 3.2 Stratifying individuals by likelihood of responding to a pre-selected dose

In the previous section, we asked the question of which set of combination doses are robustly Pareto optimal for individuals that fall within specific parameter voxels. However, for approved combinations, it may not always be possible to vary doses or schedules. To address this, we next ask the following question: for a fixed set of doses/schedule, for what individuals (i.e., for which parameter voxels) would that combination optimally balance synergy of efficacy and potency?

To answer this, we perform the following analysis: for each dose combination (as discretized in the heatmaps in Figures 3-5), we calculate the Pareto Optimality Score 𝒫 for each parameter voxel. Four representative outcomes, for four different doses of pembrolizumab plus bevacizumab, are shown in Figure; a video summary of the results across the entirety of 10×10 combination dosing space is available in the Online Supplementary Materials. All results are still shown for a TGI cutoff of at least 60%, i.e., the plots show the Pareto Optimality Score 𝒫 for each voxel provided the combination for that voxel achieves at least 60% tumor growth inhibition relative to the control. If 60% inhibition is not achieved, nothing is visualized for that parameterization.

Figure demonstrates that individuals with different drug-response characteristics respond very differently to the same combination dose. For instance, for the fixed dose of P2.7/B0.1 (Figure A), the ideal subgroup is either a strong responder to bevacizumab monotherapy (high *k*2_*d*2_)) or the combination therapy (high *k*_3_). However, they cannot be strong responders to pembrolizumab (lower k2_*d*1_). The six groups for which this protocol is robustly Pareto optimal (𝒫 ≥ 0.9) are: minimal/moderate/strong, mild/strong/moderate, mild/strong/strong, minimal/strong/strong, moderate/strong/strong, and mild/strong/mild.

As another example, for the fixed dose of P3.9/B0.5, the ideal subgroup must be a strong responder to both monotherapies but weak responders to the combination therapy (Figure B). There are only two subgroups for which this protocol is robustly Pareto optimal: strong/strong/minimal and strong/strong/mild. If we next consider the fixed dose of P2.7/B0.9, the ideal individual must be a moderate to strong responder to pembrolizumab monotherapy (Figure C). Though, our confidence in this prediction is somewhat lower than the prior doses, as no subgroup has 𝒫 ≥0.9. The top three subgroups have and correspond to the following classifications: strong/moderate/mild, strong/moderate/minimal, and moderate/minimal/moderate.

While these examples are indicative of how to identify optimal subgroups for a specific protocol, Figure D shows that there is no guarantee that a robustly optimal subgroup exists for each protocol. For the high doses of P6.2/B3.5 (Figure D), Pareto optimality scores are minimal across all voxels, with never exceeding 0.25 across all subgroups. This strongly suggests that this protocol is a poor choice independent of the patient population.

Taken together, these results highlight that knowing where an individual’s parametrization falls within the range of possibilities can be critical for finding a robustly Pareto optimal combination. It also shows, intriguingly, that lack of response may stem not from the poorly chosen combination, but from poorly selected doses. Furthermore, there does not appear to be an intuitive “rule” that can predict how the interplay of these drug sensitivities will be realized with respect to selecting doses that are more likely to balance synergy of efficacy and potency. A possible approach to identify these parameters will be discussed next.

### 3.3 Individual classification using an initiation protocol

In the analysis presented so far, we demonstrated how combination dose(s) that are Pareto optimal for any individual voxel can be identified and how to find the individual voxel(s) that best balance synergy of potency and efficacy for a pre-selected combination dose. However, in the absence of measurable biomarkers, it is unclear how one can identify, for each specific individual, to which voxel they belong. That is, since each voxel is determined by a narrow range of three parameters that capture one’s response to both monotherapy and combination therapy, it is unclear how one can measure these characteristics *a priori* to allow for this type of stratification.

To address this critical question, we propose administration of an “initiation protocol”, which could feasibly be administered to each individual at the beginning of the treatment to determine their individual parameter values. For proof of concept, we propose the following protocol:

1. On day 0, measure tumor volume and administer the drug with the shortest half-life. Here, it is bevacizumab, administered at the experimental dose of 1 mg/kg. Wait for the drug to clear before administering the next dose and again measuring the tumor volume. Here we define “clearance” as when the drug plasma levels reduce to 90% below Cmax, although other relevant criteria that meet this objective, such as reaching the lower limit of quantification (LLOQ) could be used. With all other parameters fixed, this allows us to determine a unique value of *k*2_*d*2_. It should be noted that either drug can be administered first but the one with a shorter half-life (here bevacizumab) will allow for a shorter duration of the pre-initiation protocol (24 days to meet the clearance criterion versus 31 days for pembrolizumab).
2. Pembrolizumab is then administered at the experimental dose of 10 mg/kg after the first drug has been eliminated, here on day 24. After the administration of pembrolizumab, we wait six days, enough time for pembrolizumab levels in the plasma to surpass its Cmax, and measure the tumor volume. Cmax is chosen as a reproducible criterion; the key consideration is that the second drug needs to have had the time to have measurable effect. With all other parameters fixed, this allows us to determine a unique value of *k*2 _d1_.
3. On day 30, the same day the last measurement was taken, the combination dose is administered. Note that at this stage, there was no need to wait for pembrolizumab to be effectively cleared to administer the combination dose, as having pembrolizumab present does not confound the computation of the combination effect once the other parameters have been estimated. We wait for 6 more days, again allowing both drugs to surpass their peak plasma concentration level, to measure tumor volume. Given that we already know and , this allows us to uniquely determine.

A summary of the initiation protocol, along with sample output for a representative individual, is given in Figure. More details are provided in Supplementary Figure 4. Notably, each therapy-response parameter is practically identifiable (Raue et al. 2009; Eisenberg & Jain 2017) for this initiation protocol, as illustrated by the sharp increase in the value of the cost function as each parameter deviates from its optimal value. The figure also provides sample output of the algorithm with a summary of the top combination doses, in terms of Pareto optimality, for this representative individual. We define the “top” doses as all combination doses with a score within 25% of the combination with the largest.

This approach allows identifying individual parameters to place them in the appropriate voxel, which then permits either identifying a protocol that best balances synergy of efficacy and potency, or assessing whether they are a good candidate to benefit from a pre-established protocol.

## 4 DISCUSSION

The process of drug development does not end with identification of a promising compound, determining its safety and efficacy, and getting it to the clinic. Patient stratification can determine whether a compound will be successful or not. This process can be difficult even for monotherapies and is particularly challenging for combination therapies. Identifying characteristics of those individuals for whom the dose is expected to work is particularly important, since lack of efficacy in a poorly chosen subset of individuals may result in the compound or a combination being pulled because the wrong patient population was considered.

In this work, we propose a methodology for treatment semi-personalization based on the simultaneous optimization of synergy of efficacy and synergy of potency (a proxy for treatment toxicity). This work expands upon our previously developed MOOCS-DS framework for identifying Pareto optimal combination therapies (Gevertz & Kareva 2023) across various definitions of synergy. Here, a dose is Pareto optimal if one measure of synergy (say, synergy of efficacy) cannot be improved without worsening the other measure of synergy (synergy of potency). As a case study, we apply the methodology to a preclinical model of the combination of an immune checkpoint inhibitor and an anti-angiogenic agent. We use the term “semi-personalized” because each individual is classified into a subgroup based on if their response to the two monotherapies and the combination is classified as either minimal, mild, moderate, or strong. This is in comparison to both a population-level study which would simply determine treatment response using the average drug-response parameters, and a personalized approach, which would require the precise value of each drug-response parameter for each individual. For each treatment-response subgroup, which we call a parameter voxel, we compute the Pareto Optimality Score 𝒫 as a measure of the robustness of an optimality prediction across various definitions of synergy and across all individuals that fall within the subgroup. This approach thus provides a quantifiable assessment for why the same combination doses well-balance efficacy and toxicity for some subsets of individuals but not for others.

Using the Pareto Optimality Score as a criterion for semi-personalized dose selection, we first ask the following question: if one knows which parameter voxel an individual falls into, what would be the most robust Pareto optimal dose for those individuals? In answering this question across parameter voxels, we find that the most robust Pareto optimal dose(s) can vary dramatically depending on the drug-response characteristics of a subgroup. We also find that there are subgroups that have quite a homogeneous response, making the choice of a robust Pareto optimal combination therapy clear. However, there are also examples where there is quite a lot of heterogeneity within a subgroup, which makes finding a Pareto optimal dose for individuals with those characteristics particularly challenging.

We next asked the following question: if a combination dose is pre-determined (i.e., from a pre-established protocol), for which subgroups would this combination dose provide a robust Pareto optimal solution? Our methodology not only provides an answer to this question but also reveals that there is not a simple intuitive “rule” for predicting how the interplay between drug sensitivities will be realized with respect to selecting robust Pareto optimal doses that simultaneously optimize synergy of efficacy and potency.

Finally, it is critically important to identify to which voxel an individual belongs. Ideally, one would use a validated measurable biomarker, such as PSA for prostate cancer (Catalona 2014), or soluble BCMA for multiple myeloma (Lee et al. 2024). These biomarkers are not flawless, but they could facilitate patient stratification (Brady-Nicholls et al. 2021; Gevertz & Kareva 2025). In some cases, a genetic biomarker may be feasible, such as staining for estrogen receptor (ER+). However, there do exist challenges with this type of marker, for instance when there is a mismatch between treatment inclusion criteria (1% ER+ cells, (Hammond et al. 2010)) and anticipated efficacy (requiring over 10% ER+ (Morgan et al. 2011)). Additional complications arise from variability in classification, which can depend on the region of the heterogeneous tumor selected for biopsy, as well as inter-lab variability (Layfield et al. 2003).

Unfortunately, not all types of cancer have validated measurable biomarkers, and a particular challenge arises when it is known that a “rate” parameter is going to be critical (i.e., rate at which a drug kills cancer cells), but that rate cannot be directly measured. In such cases, one can conceivably only measure the outcome of the effect and then infer the what the rate must have been to achieve such an outcome. For these scenarios, we propose an “initiation protocol”, which allows in a reproducible and structured way to infer what the rate of response must have been to result in a particular outcome. This initiation protocol must be carefully designed so that the data collected allows for the inference of a unique value for the parameter; that is, the parameter of interest must be practically identifiable given the data collected in the initiation protocol.

The implementation of our proposed initiation protocol allows individuals to be appropriately stratified. This allows us to determine the Pareto optimal doses for their specific voxel, to assess whether they are a candidate for a pre-selected dose, or to determine that this combination of drugs does not result in a robust Pareto optimal solution with sufficient TGI. Notably, we propose using voxels as opposed to individualized therapy recommendations because, as mentioned above, there may exist variability in data collection and measurements. By relying on voxel-level groupings of parameter values instead of exact individualized measurements, our approach introduces a buffer that helps ensure prediction robustness, even when parameter estimates are uncertain or imprecise.

While we have sought to show the viability of our proposed approach, there exist limitations and caveats that need to be understood and addressed before its realistic implementation. First, in our preclinical example, even though murine pembrolizumab has a half-life of only 2-3 days (Lindauer et al. 2017), it still takes over three weeks for a dose of 10 mg/kg to clear out to under 90% of Cmax. It is important for this to occur before the initiation dose of the second drug is administered to be able to isolate the relative contributions to efficacy of one monotherapy compared to the other. Over three weeks of time, however, may not be realistic for a mouse experiment, where animals may need to be sacrificed for ethical reasons at earlier time points if the tumor grows to be too large. One alternative approach is to use a lower dose, which will reach the LLOQ sooner (without impacting the criterion of waiting for concentration to drop below 90% of Cmax). However, one might not be able to correctly estimate the parameter if the dose is subtherapeutic. Furthermore, it is possible that the mechanism of action of the drug may have a delay, i.e., if a drug’s effect shows hysteresis, in which case a single dose may not be sufficient to properly estimate the drug kill parameter. As such, the applicability of this approach may need to be tailored based on both the drug half-life and the anticipated mechanism of action.

Ideally, the proposed initiation protocol could be complemented by in vitro experiments, such classical co-incubation experiments (Mason & Öhlund 2023), to approximate values of individual-specific parameters of sensitivity to treatment. One critical drawback of such an approach is lack of the input of the tumor microenvironment, which can be critical for how efficacious the drug may actually be in an individual; a 3D tissue organoid may be a better proxy for such an ex vivo approach (Tosca et al. 2023) and remains to be evaluated. The notion of co-clinical trials, where a tumor xenograft from an individual patient is concurrently grown in a mouse (Kim et al. 2020), is another option to approximate these drug-response parameters. It remains to be evaluated whether such an approach would allow estimating the parameters well enough to classify an individual into the appropriate parameter voxel.

Second, the methodology proposed requires having some assessment of efficacy subject to the drug. In a preclinical setting, this data is readily accessible using calipers to measure tumor volume. However, in a clinical setting, such well-resolved time course data is rarely available, with the exception of blood cancers. This poses challenges for translating the proposed methodology from a preclinical to a clinical setting. In this clinical setting, one either has to rely on imaging data that is often limited by accessibility and/or insurance coverage, the availability of a validated biomarker of response, or potentially relative reductions in circulating tumor DNA for cancers that have a sufficiently high rate of DNA shedding (Sanchez-Herrero et al. 2022). Translation of the method to clinical data will thus be context specific, depending on the data available.

Third, the case study presented herein is performing uncertainty quantification on the three model parameters that describe drug sensitivity. There are other forms of uncertainty that are not assessed under this framework: uncertainty in the choice of model type (e.g., ordinary differential equations, partial differential equations, agent based), uncertainty in the functional forms used in the PK and PD equations, and uncertainty in the parameters that were fixed in order to form the subgroups. Regarding the latter, the proposed methodology does generalize beyond three-dimensional parameter space and could incorporate uncertainty in as many parameters as deemed necessary. In the context of a specific clinical application, care will be required to determine which model parameters are used for patient subgrouping. This case study uses three drug sensitivity parameters to illustrate how our methodology captures the way inter-individual variability necessitates distinct, optimally synergistic dose predictions for each subgroup.

The method will also confront other challenges when being applied to clinical data. Preclinical studies are conducted over short time scales, and thus it is reasonable to assume constant treatment response parameters over the course of the study. However, in a clinical setting, it cannot be ignored that treatment-naïve tumor may have different sensitivity parameters than the same tumor even after a few treatment cycles. The possibility of evolving resistance is not yet considered in the proposed framework and is something that must be accounted for in future work that extends the method to work for clinical data. Similarly, incorporating variability in the pharmacokinetic parameters, as well as more mechanistically detailed PK models that account for nonlinear clearance mechanisms such as target-mediated drug disposition, represents an important direction for future work, particularly as this framework is extended towards clinical application.

In summary, it is critical to account for inter-individual variability in sensitivity to therapeutics to ensure that promising therapies are not discarded prematurely due inadequate stratification. The methodology proposed herein represents a step in how we can one day ensure that we deliver the right combinations to the right patients, and that we identify the right patients for specific combinations.

## ACKNOWLEDGEMENTS

JLG acknowledges use of the ELSA high-performance computing cluster at The College of New Jersey for conducting the research reported in this paper. This cluster is funded in part by the National Science Foundation under grant numbers OAC-1826915 and OAC-1828163. JLG and IK are grateful to Dr. Kathleen Wilkie for her suggestion to consider near Pareto optimal solutions, and to Kyla Devlin for her work in expanding the MOOCS-DS algorithm to find these near Pareto optimal points. JLG and IK also wish to thank Dr. Victoria Prince for very helpful discussions on methodological approaches to data analysis. The authors would like to dedicate this work to the late Dr. Siv Sivaloganathan, who brought us all together as a community both in Fields Institute and beyond. JLG appreciated sharing the original MOOCS-DS algorithm with him at the Frontiers in Computation and Mathematical Medicine Workshop in September of 2024, and for how Siv welcomed her into his academic family during that workshop.

## AUTHOR CONTRIBUTIONS

Both authors contributed equally to writing the manuscript, designing research, performing research, and analyzing data.

## CODE AVAILABILITY

All code is available on GitHub at https://github.com/jgevertz/MOOCS-DS/tree/main/Variability.

## SUPPLEMENTARY MATERIALS

### S1. Supplementary Methods: Details of Original MOOCS-DS Algorithm

Synergy is defined relative to additivity, where if a combined effect is greater than what is expected in the additive case, it is classified as synergistic, and when it is less than expected from additivity, it is classified as antagonistic. Additivity definitions can broadly be divided into two categories: those focused on the efficacy of treatment (synergy of efficacy, SoE) and those focused on the input of the treatment (synergy of potency, SoP).

As the names suggest, SoE metrics assess whether the output of the combination (the efficacy) is greater than the one that would have been expected from an additive case. Bliss additivity, which defines additivity via the notion of drug independence, is the most commonly used measure for SoE (Bliss 1939). For Bliss additivity, define *E* as the normalized efficacy (between 0 and 1) of a combination therapy that administers a dose *d*_1_ of drug 1 and *d*_2_ of drug 2. Additivity is defined as achieving an efficacy of:

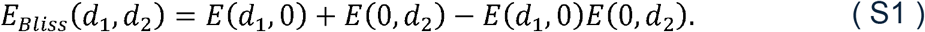

If the efficacy of the combination exceeds *E*_Bliss_ (*d*_1_,*d*_2_) , the combination is classified as synergistic. As Bliss is known to set a very high bar for additivity, other SoE metrics have been proposed. Another common one is Highest Single Agent (HSA), which defines additivity as the combination having an efficacy equal to the more efficacious monotherapy (Vlot et al. 2019):

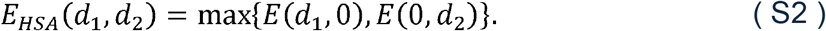

HSA sets a very low bar for additivity, as synergy is achieved with any improvement in efficacy beyond the more effective monotherapy. To facilitate the comparison of synergy scores across additivity metrics, the combination index (*CI*) is used (Chou & Talalay 1983). For SoE, this is defined as the ratio of the expected additive efficacy to the actual combination efficacy:

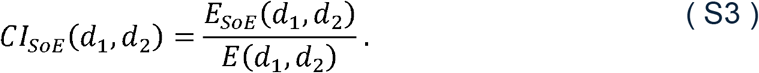

Using this metric, a combination is additive if its *CI* equals 1. Values greater than 1 represent antagonism, with larger values being more antagonistic. Values less than 1 represent synergy, with smaller values being more synergistic.

SoP metrics assess how much drug must be administered to achieve a fixed efficacy, with drug concentrations less than additive being classified as synergistic. The most equivalence (Loewe 1926). In particular, assume there is a combination dose common SoP metric is Loewe, where additivity is defined by assumption of dose (*d*_1_,*d*_2_) that achieves the same effect as the monotherapy of drug 1 at dose *D*_1_ and drug 2 at dose *D*_2_:

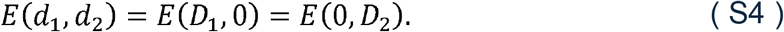

Loewe’s definition of additivity is given by

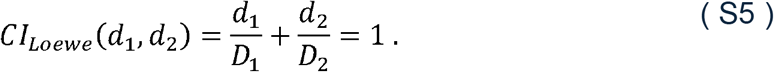

As with SoE metrics, *CI* values below 1 represent synergy (less drug is needed than predicted by monotherapy response to achieve a fixed efficacy) and values above 1 represent antagonism. The *CI* equation for Loewe additivity is derived assuming the two drugs have parallel dose-response curves, which does limit its applicability in cases where this assumption is far from holding.

In (Gevertz & Kareva 2023) we also introduced the Lowest Single Dose (LSD) metric as a potency-based equivalent to HSA. LSD classifies a dose as additive if potency of the combination is equal to that of the more potent monotherapy. Like Loewe, LSD considers the case of a combination dose (*d*_1_,*d*_2_) that achieves the same efficacy as the monotherapies (eqn. (S4)) and defines the *CI* as:

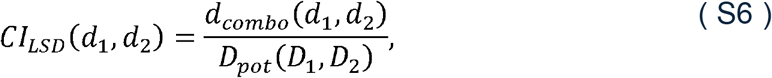

where *d*_*combo*_ (*d*_1_,*d*_2_) is a normalized measure of the combination dose and *D*_*pot*_(*D*1,*D*2) is a normalized measure of the more potent monotherapy. To detail, define the 50% inhibition value (*PI*_50_) as the dose of a drug that results in 50% tumor growth inhibition relative to the control. Then:

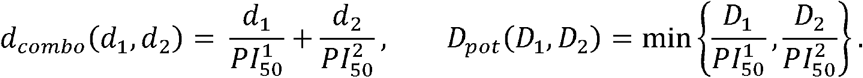

As with all additivity definitions, *CI*_*LSD*_ = 1 represents additivity, *CI*_*LSD*_ < 1 represents synergy, and *CI*_*LSD*_ > 1 represents antagonism.

While each of these definitions have their own merits, drawing meaningful conclusions using any single definition is ill-advised since different definitions result in different classifications (Vlot et al. 2019). To address this inconsistency, in (Gevertz & Kareva 2023) we introduced a Pareto optimization approach that identifies combination doses for which one objective (i.e., synergy of potency) cannot be improved without the other (i.e., synergy of efficacy) worsening. The set of doses that satisfy this tradeoff is called the Pareto front. These combination doses should be thought of as optimally balancing treatment effectiveness (measured through SoE) and toxicity (measured through SoP).

### S2. Supplementary Figures and Tables

**Supplementary Figure S1.**
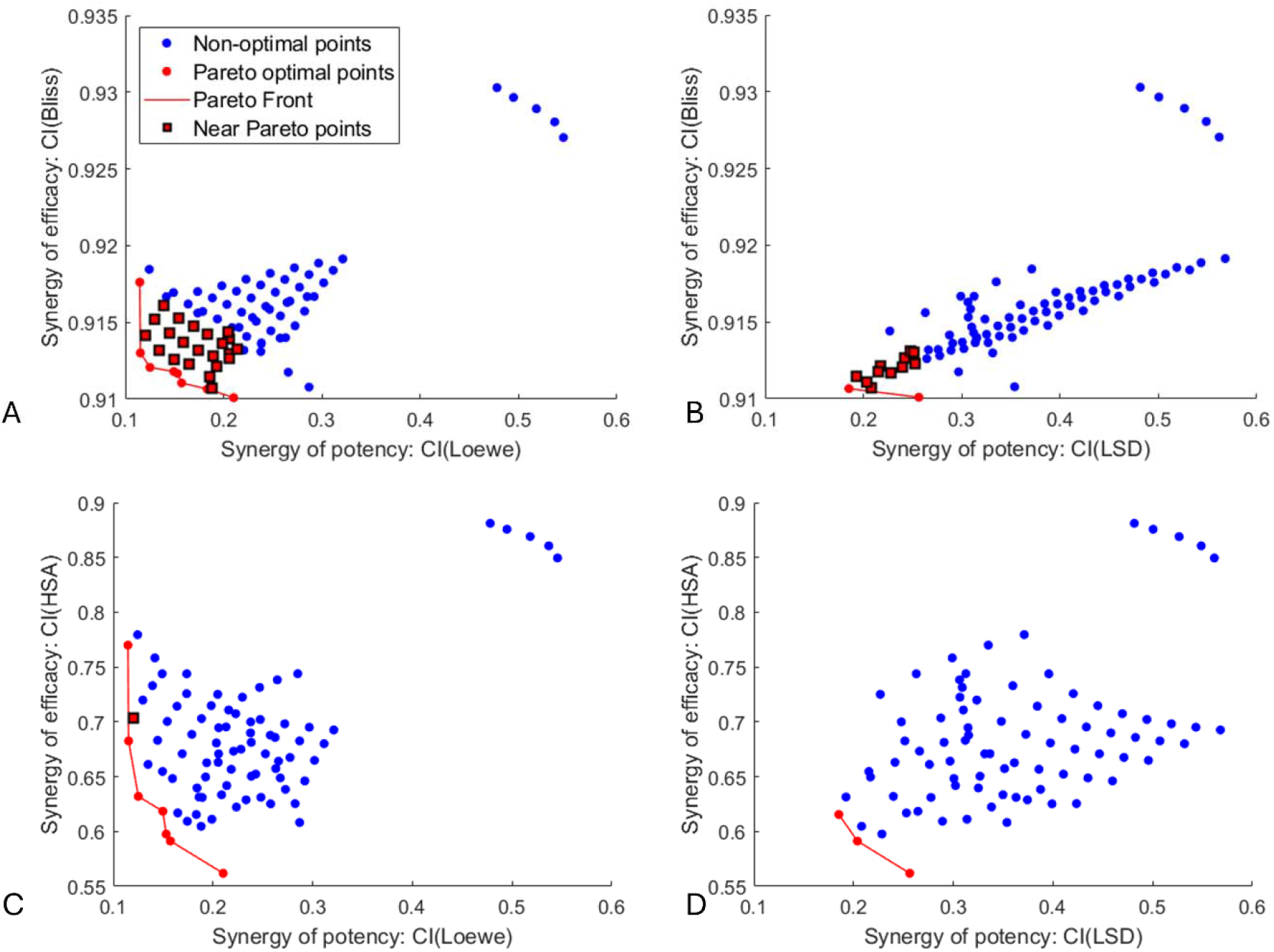
Pareto optimal solutions (red circles), near Pareto points (red squares outlined in black), and non optimal points (blue circles) in (A) Loewe-Bliss space, (B) LSD-Bliss space, (C) Loewe-HSA space, (D) LSD-HSA space. Results are for the model in eqns. (1) – (3) solved at the nominal parameter values defined in Table 1.

**Supplementary Figure S2.**
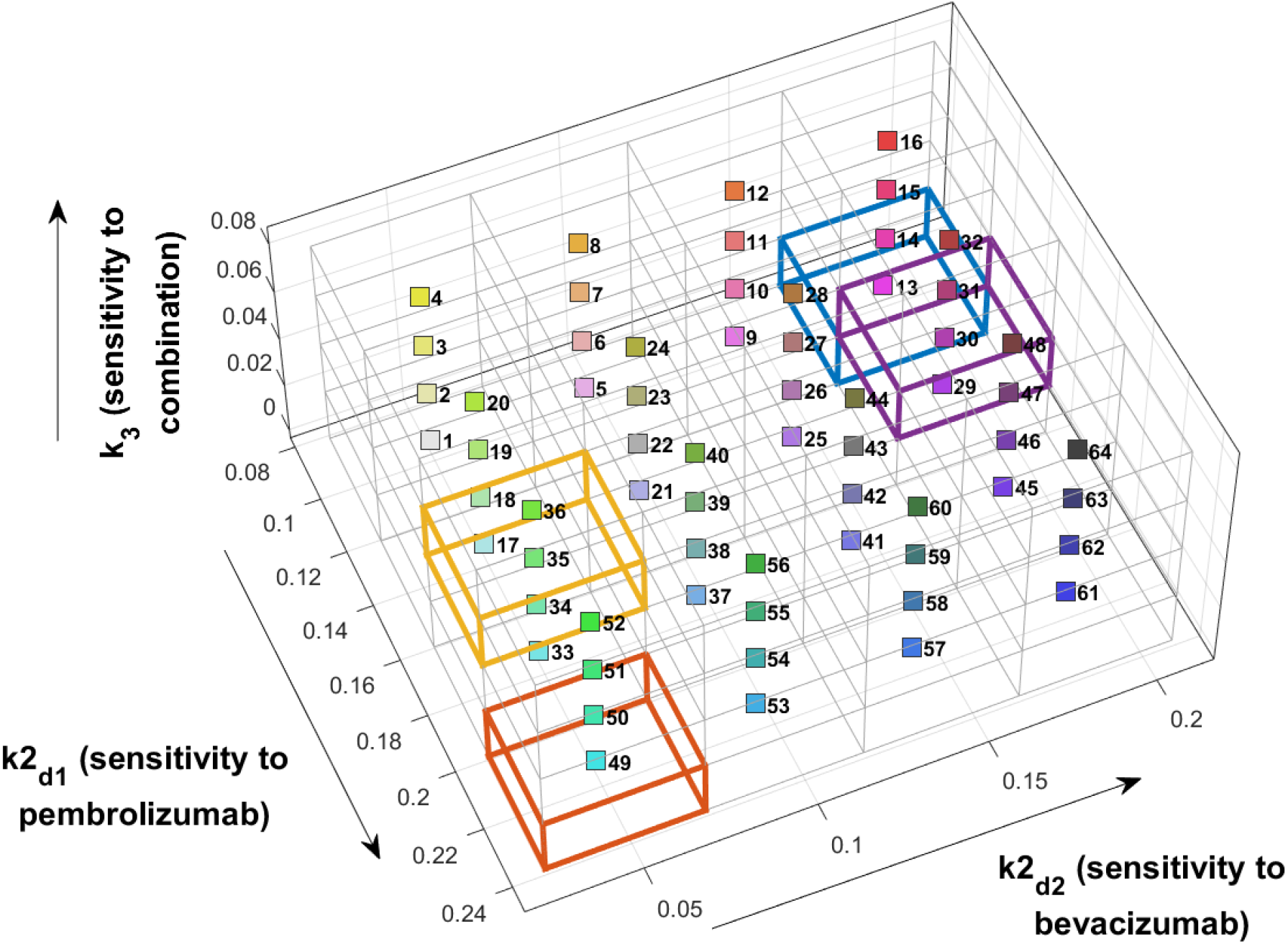
Position of voxels in the parameter space, highlighting representative voxels 13 (blue; minimal/strong/minimal response to pembrolizumab, bevacizumab and combination, respectively), 30 (purple; mild/strong/mild); 35 (yellow; moderate/minimal/moderate) and 49 (red; strong/minimal/minimal).

**Supplementary Table S1.**
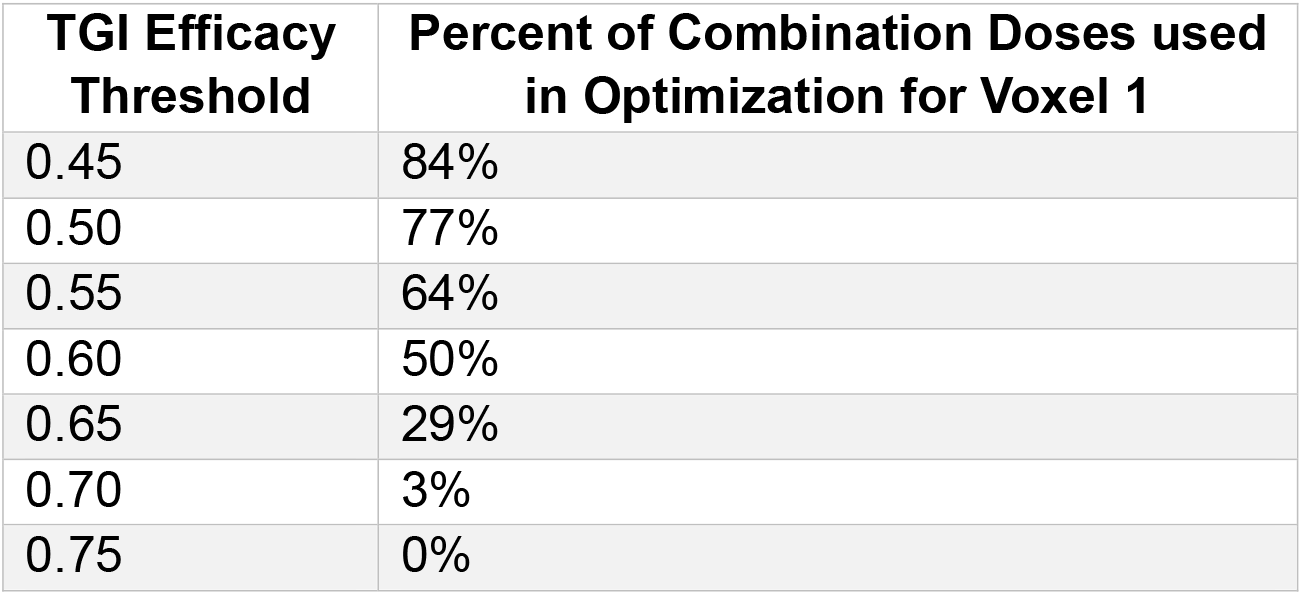
Size of feasible set for the weakest responders (Voxel 1) as a function of the TGI efficacy threshold.

| **TGI Efficacy Threshold** | **Percent of Combination Doses used in Optimization for Voxel 1** |
| --- | --- |
| 0.45 | 84% |
| 0.50 | 77% |
| 0.55 | 64% |
| 0.60 | 50% |
| 0.65 | 29% |
| 0.70 | 3% |
| 0.75 | 0% |

**Supplementary Figure S3.**
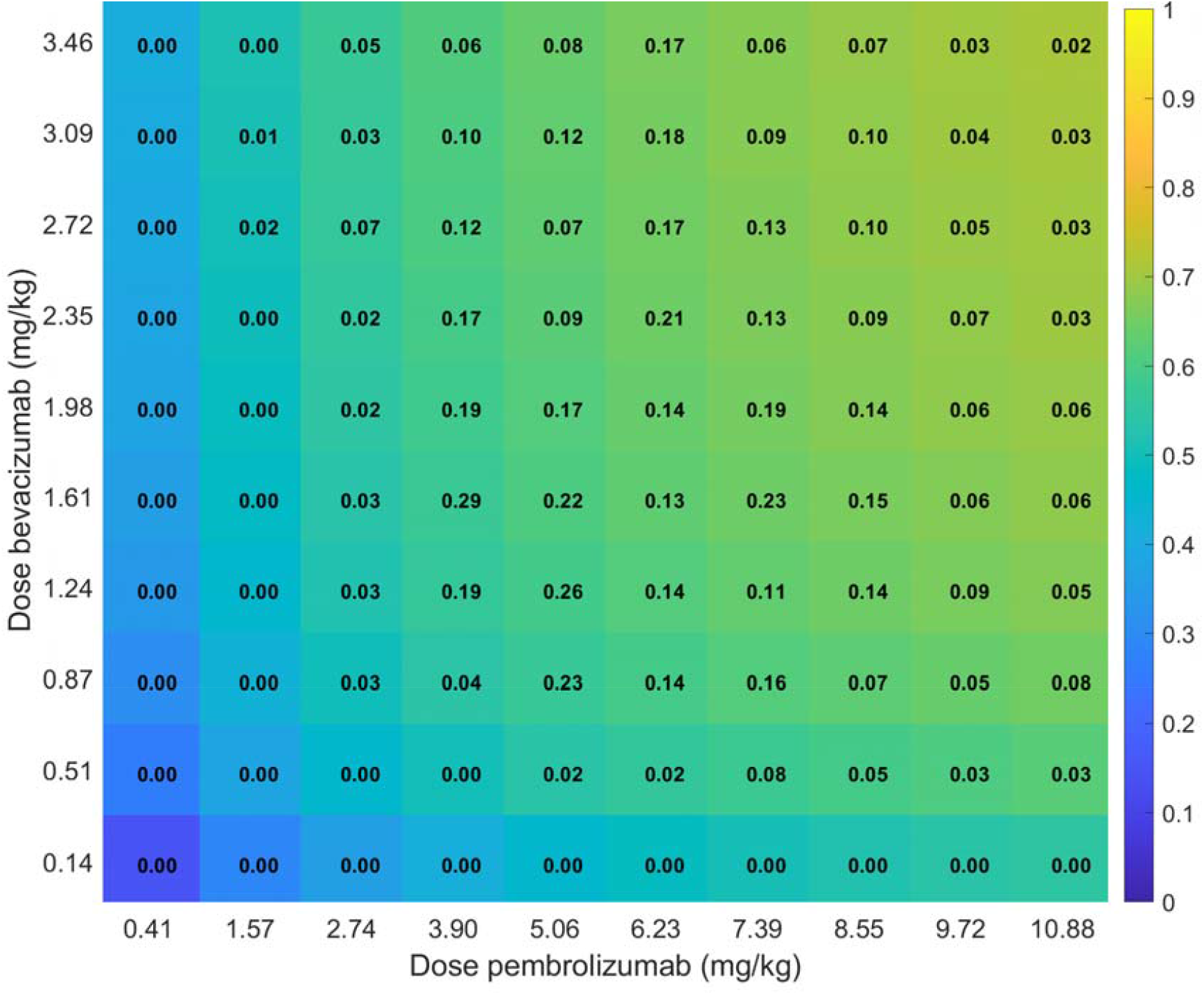
Color in the heatmap represents TGI achieved at the center of parameter voxel 1 (*k*2_*d*1_ ∈ [0.084,0.123], *k*2_d2_ ∈ [0.025,0.071], k_3_ ∈ [0.0006,0.021]). The largest TGI of 0.71 is achieved at the maximum combination dose considered. The number within each cell is the Pareto Optimality Score for the combination dose. Note Pareto optimality scores were computed using a TGI threshold of 0.6.

**Supplementary Figure S4.**
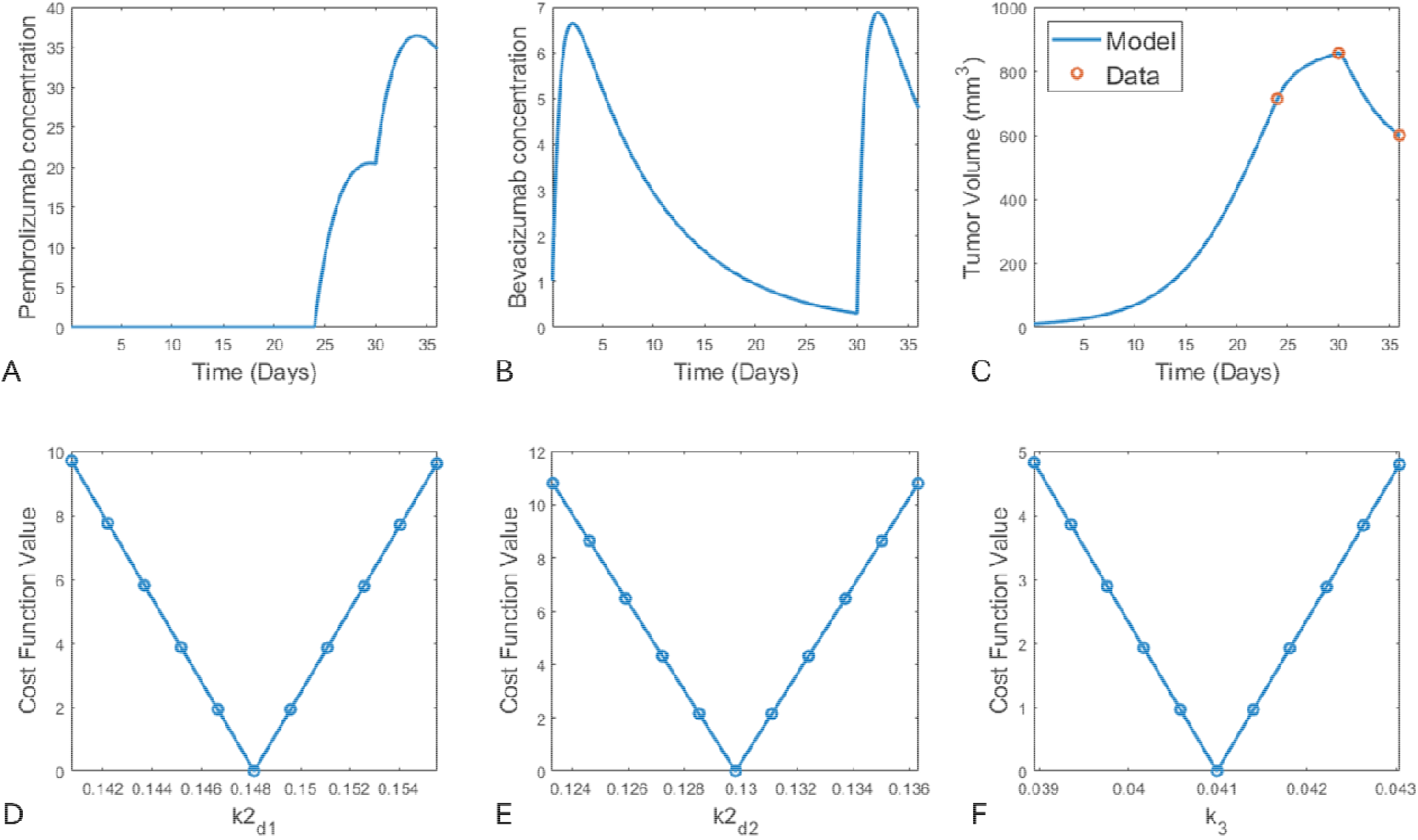
Details of initiation protocol where a dose of 1 mg/kg of bevacizumab is adminstered on Day 0, 10 mg/kg of pembrolizumab is administered on Day 24, and the combination is administered on Day 30. Output is shown for a mouse in which the tumor grows by 7000% from Day 0 to Day 24 (in response to the monotherapy of bevacizumab), grows by 20% from Day 24 to Day 30 (in response to the monotherapy of pembrolizumab), and shrinks by 30% from Day 30 to Day 36 (in response to the combination). (A) Pembrolizumab concentration over time. (B) Bevacizumab concentration over time. (C) Tumor volume time-course that results from fitting drug-response parameters, one at a time, to each data point. (D)-(F) Profile likelihood curves for drug-response parameters , , , respectively.

### S3. Online Supplementary Materials

Available at https://github.com/jgevertz/MOOCS-DS/tree/main/Variability

**Supplementary Video 1**. Pareto Optimality Score for all 64 voxels. Four representative cases (voxels 13, 30, 35 and 49) are shown in Figure 5 of the main text.

**Supplementary Video 2**. Pareto Optimality Score for all parameter voxels for all combination doses considered. Four representative doses (P2.7/B0.1; P3.9/B0.5; P2.7/B0.9 and P6.2/B3.5) are shown in Figure 6 of the main text. Parameterizations that did not meet the efficacy threshold (TGI ≥0.6) have no value shown.

**Supplementary File 1**. Values of , , that define each parameter voxel.

## REFERENCES

Avanzini, S. et al., 2020. A mathematical model of ctDNA shedding predicts tumor detection size. Science advances, 6(50), p.eabc4308.

Bliss, C.I., 1939. The toxicity of poisons applied jointly 1. Annals of applied biology, 26(3), pp.585–615.

Boakye, D. et al., 2021. Early discontinuation and dose reduction of adjuvant chemotherapy in stage III colon cancer patients. Therapeutic Advances in Medical Oncology, 13, p.17588359211006348.

Brady-Nicholls, R. et al., 2021. Predicting patient-specific response to adaptive therapy in metastatic castration-resistant prostate cancer using prostate-specific antigen dynamics. Neoplasia, 23(9), pp.851–858.

Catalona, W.J., 2014. History of the discovery and clinical translation of prostate-specific antigen. Asian Journal of Urology, 1(1), pp.12–14.

Chapman, P.B. et al., 2017. Vemurafenib in patients with BRAFV600 mutation-positive metastatic melanoma: final overall survival results of the randomized BRIM-3 study. Annals of Oncology, 28(10), pp.2581–2587.

Chou, T.-C. & Talalay, P., 1983. Analysis of combined drug effects: a new look at a very old problem. Trends in pharmacological sciences, 4, pp.450–454.

Costa, D.B. et al., 2007. Pooled analysis of the prospective trials of gefitinib monotherapy for EGFR-mutant non-small cell lung cancers. Lung cancer, 58(1), pp.95–103.

Domchek, S.M. et al., 2016. Efficacy and safety of olaparib monotherapy in germline BRCA1/2 mutation carriers with advanced ovarian cancer and three or more lines of prior therapy. Gynecologic oncology, 140(2), pp.199–203.

Eisenberg, M.C. & Jain, H.V., 2017. A confidence building exercise in data and identifiability: Modeling cancer chemotherapy as a case study. Journal of theoretical biology, 431, pp.63–78.

Gevertz, J.L. & Kareva, I., 2023. Guiding model-driven combination dose selection using multi-objective synergy optimization. CPT: Pharmacometrics & Systems Pharmacology.

Gevertz, J.L. & Kareva, I., 2025. Using Virtual Patients to Evaluate Dosing Strategies for Bispecifics with a Bell-Shaped Efficacy Curve. bioRxiv, pp.2025–5.

Hammond, M. et al., 2010. Pathologists’ guideline recommendations for immunohistochemical testing of estrogen and progesterone receptors in breast cancer. Breast Care, 5(3), pp.185–7.

Karachaliou, N. et al., 2019. EGFR first-and second-generation TKIs—there is still place for them in EGFR-mutant NSCLC patients. Translational Cancer Research, 8(Suppl 1), p.S23.

Kazandjian, D. et al., 2016. FDA approval of gefitinib for the treatment of patients with metastatic EGFR mutation–positive non–small cell lung cancer. Clinical Cancer Research, 22(6), pp.1307–1312.

Kent, D.M. et al., 2020. The predictive approaches to treatment effect heterogeneity (PATH) statement. Annals of internal medicine, 172(1), pp.35–45.

Kim, H.R. et al., 2020. Mouse–human co-clinical trials demonstrate superior anti-tumour effects of buparlisib (BKM120) and cetuximab combination in squamous cell carcinoma of head and neck. British journal of cancer, 123(12), pp.1720–1729.

Layfield, L.J. et al., 2003. Interlaboratory variation in results from immunohistochemical assessment of estrogen receptor status. Breast Journal, 9(3).

Lee, H. et al., 2024. Impact of soluble BCMA and non-T cell factors on refractoriness to BCMA targeting T cell engagers in multiple myeloma. Blood.

Lin, Y.S. et al., 1999. Preclinical pharmacokinetics, interspecies scaling, and tissue distribution of a humanized monoclonal antibody against vascular endothelial growth factor. Journal of Pharmacology and Experimental Therapeutics, 288(1), pp.371–378.

Lindauer, A. et al., 2017. Translational pharmacokinetic/pharmacodynamic modeling of tumor growth inhibition supports dose-range selection of the anti–PD-1 antibody pembrolizumab. CPT: pharmacometrics & systems pharmacology, 6(1), pp.11–20.

Loewe, S., 1926. Effect of combinations: mathematical basis of problem. Arch. Exp. Pathol. Pharmakol., 114, pp.313–326.

Luo, M.C., Nikolopoulou, E. & Gevertz, J.L., 2022. From fitting the average to fitting the individual: A cautionary tale for mathematical modelers. Frontiers in Oncology, p.1311.

Mason, J. & Öhlund, D., 2023. Key aspects for conception and construction of co-culture models of tumor-stroma interactions. Frontiers in Bioengineering and Biotechnology, 11, p.1150764.

Morgan, D.A., Refalo, N.A. & Cheung, K.L., 2011. Strength of ER-positivity in relation to survival in ER-positive breast cancer treated by adjuvant tamoxifen as sole systemic therapy. The Breast, 20(3), pp.215–219.

Nielson, C.M. et al., 2021. Relative dose intensity of chemotherapy and survival in patients with advanced stage solid tumor cancer: a systematic review and meta-analysis. The Oncologist, 26(9), pp.e1609–e1618.

Qiao, T. et al., 2023. Combined pembrolizumab and bevacizumab therapy effectively inhibits non-small-cell lung cancer growth and prevents postoperative recurrence and metastasis in humanized mouse model. Cancer Immunology, Immunotherapy, 72(5), pp.1169–1181.

Raue, A. et al., 2009. Structural and practical identifiability analysis of partially observed dynamical models by exploiting the profile likelihood. Bioinformatics, 25(15), pp.1923–1929.

Sanchez-Herrero, E. et al., 2022. Circulating tumor DNA as a cancer biomarker: an overview of biological features and factors that may impact on ctDNA analysis. Frontiers in Oncology, 12, p.943253.

Scibilia, K.R. et al., 2025. Mathematical Oncology: How Modeling Is Transforming Clinical Decision Making. Cancer research.

Shin, S. et al., 2024. Discontinuation risk from adverse events: immunotherapy alone vs. combined with chemotherapy: a systematic review and network meta-analysis. BMC cancer, 24(1), p.152.

Spreafico, M., Ieva, F. & Fiocco, M., 2024. Causal effect of chemotherapy received dose intensity on survival outcome: a retrospective study in osteosarcoma. BMC Medical Research Methodology, 24(1), p.296.

Tanniou, J. et al., 2016. Subgroup analyses in confirmatory clinical trials: time to be specific about their purposes. BMC medical research methodology, 16(1), p.20.

Tosca, E.M. et al., 2023. Replacement, reduction, and refinement of animal experiments in anticancer drug development: the contribution of 3D in vitro cancer models in the drug efficacy assessment. Biomedicines, 11(4), p.1058.

Vlot, A.H. et al., 2019. Applying synergy metrics to combination screening data: agreements, disagreements and pitfalls. Drug discovery today, 24(12), pp.2286–2298.

Xia, X. et al., 2023. The history and development of HER2 inhibitors. Pharmaceuticals, 16(10), p.1450.

